# Differential Branched-Chain Amino Acid Metabolism in Tissues of Tumor-Bearing Mice

**DOI:** 10.1101/2025.11.01.685925

**Authors:** Miriam Zerrouk, Luca J. Delfinis, Christopher G.R. Perry, Olasunkanmi A.J Adegoke

## Abstract

Cancer cachexia is a multifactorial syndrome characterized by the involuntary loss of skeletal muscle and adipose tissue, often resistant to nutritional support. The branched-chain amino acids (BCAA: leucine, isoleucine, and valine) stimulate protein synthesis, yet BCAA-targeted therapies have yielded limited clinical benefit, and inconsistent results. In this study, a C26 colon cancer mouse model was used to examine how tumor burden alters BCAA metabolism across skeletal muscle, liver, kidney, and adipose tissue. Tumors accumulated BCAA and showed increased oxidation of these amino acids, whereas peripheral sites displayed widespread BCAA depletion, reduced expression of the amino acid (AA) transporter LAT1, and suppression of mechanistic target of rapamycin complex 1 (mTORC1) signaling. Notably, the soleus muscle maintained mTORC1 activity despite reduced BCAA availability, suggesting fiber-type–specific adaptations. These findings indicate that tumors act as metabolic sinks, diverting systemic AA away from host tissues. Such reprogramming may underlie the limited success of BCAA-based interventions in cachexia and highlight the need for therapies that address both tumor and host metabolism.

**New and Noteworthy:** This is the first study to profile branched-chain α-keto acid (BCKA) levels together with branched-chain amino acid (BCAA) metabolism across multiple tissues in a cancer cachexia model. Tumors accumulated BCAA while some peripheral sites showed depletion, and all peripheral tissues exhibited reduced expression of the transporter LAT1. These tissue-specific adaptations reveal systemic metabolic reprogramming and may explain the limited efficacy of BCAA-based therapies.

## Introduction

Cancer is the second leading cause of death worldwide (1), affecting millions every year (2). Depending on the cancer type, more than half of patients experience a debilitating condition known as cachexia (3). Cancer cachexia is a devastating multifactorial metabolic syndrome, characterized by the progressive involuntary weight loss, skeletal muscle atrophy, and adipose tissue depletion (4,5) all of which reduce treatment tolerance (6), diminish quality of life, and increase mortality (7). The prevalence and severity of cachexia vary by tumor type, with pancreatic, gastrointestinal and lung cancers showing the highest rates (8). Unlike malnutrition, cachexia is not simply the result of reduced caloric intake; rather, it reflects systemic metabolic disturbances that impair anabolic processes even in the presence of nutritional support (9,10). With no effective therapy available, identifying the metabolic mechanisms that underlie tissue wasting remains essential for the development of new interventions.

The branched-chain amino acids (BCAA: leucine, isoleucine, and valine) are critical regulators of body weight and muscle mass (11,12). Leucine, in particular, activates the mechanistic target of rapamycin complex 1 (mTORC1) pathway, which promotes protein synthesis and inhibits proteolysis (13,14). Although BCAA supplementation has been tested as a potential therapeutic strategy, its effects in cancer cachexia are inconsistent, with some studies showing modest benefits and others reporting little to no impact (15). These discrepancies may be related to altered tissue metabolism and redistribution of these AAs in the cachectic state.

BCAA uptake into cells is primarily mediated by the L-type amino acid (AA) transporter 1 (LAT1) and small neutral AA transporter 1 (SNAT1), which together regulate intracellular availability for protein synthesis and signaling (16,17). In addition to their signaling roles, BCAA also serve as metabolic substrates. During catabolic states, BCAA are transaminated by branched-chain aminotransferases (BCAT1/2), generating branched-chain α-keto acids (BCKAs: 2-ketoisocaproate, 2-ketoisovalerate, and 2-keto-β-methylvalerate) and glutamate. BCKAs can be irreversibly oxidized by the branched-chain α-keto acid dehydrogenase (BCKD) complex to produce acyl-CoA derivatives that fuel the tricarboxylic acid (TCA) cycle (18,19). Beyond serving as intermediates, BCKAs are increasingly recognized as signaling molecules that influence insulin sensitivity and metabolic adaptation, particularly in conditions such as obesity and type 2 diabetes (20). Their role in cancer cachexia, however, remains largely unexplored.

In tumor cells, BCAA catabolism is often upregulated, with increased expression of BCAT and BCKD contributing to sustained AA utilization and substrate supply for tumor growth (21,22,23). This tumor demand may compete with host tissues, leading to systemic BCAA depletion and impaired anabolic signaling in muscle, liver, kidney, and adipose tissue (24,25). While previous studies have examined plasma and tumor BCAA levels in cachexia (26), less is known about how BCAA and BCKA metabolism are altered across multiple host tissues in the tumor-bearing state.

Here, we investigated BCAA concentrations, BCKA levels, BCKD activity, AA transporter expression and mTORC1 signalling in tumor, skeletal muscle, liver, kidney, and adipose tissue in a C26 mouse model of cancer cachexia. We hypothesized that cachexia would be associated with tissue-specific alterations in both BCAA and BCKA metabolism, reflecting differential AA demands across organs. To our knowledge, this is the first study to profile BCKA metabolism alongside BCAA regulation across multiple host tissues in cancer cachexia. By characterizing these changes, our study provides new insight into systemic AA regulation in cancer cachexia and may help explain the limited efficacy of current BCAA-based therapeutic strategies.

## Materials and Methods

The animal experiments and tissue collection were as described by Delfinis et al (27). Frozen tissues collected from that study were analyzed. Tissue samples were collected from n=10 for control mice (C) and n=9-10 for C26 tumor bearing mice (C26) except for the soleus muscle for which n = 5 for control and tumour bearing groups.

### Animal Care

Male CD2F1 mice (8 weeks old) were purchased from Envigo and acclimatized for a minimum of 72 hours upon arrival. During this period, mice were housed in standard laboratory conditions with ad libitum access to chow and water. Although food intake was not directly assessed, prior research with this model has shown that cachexia-induced muscle atrophy occurs independently of reduced food intake as pair-fed control mice maintained normal muscle weights, fiber cross-sectional areas, and muscle strength, in contrast to cachectic mice (28, 29). Mice were monitored daily for overall health, tumor size, and signs of distress (27).

### C26 Cell Culture and Tumor Implantation

C26 cancer cells (purchased from NIH National Cancer Institute) were cultured in T-75 flasks using Dulbecco’s Modified Eagle Medium (DMEM) supplemented with 10% fetal bovine serum (FBS) and 1% penicillin-streptomycin. Cells were maintained at passages 2–3, and once confluent, they were trypsinized, counted, and suspended in sterile phosphate-buffered saline (PBS). C26 cells (5 × 10) in 100 μL of PBS were subcutaneously injected into the flanks of experimental mice at 8 weeks of age. Control mice received identical subcutaneous injections of 100 μL of PBS and were aged for two to four weeks. Tumors were allowed to grow for 26–29 days, during which tumor dimensions were measured daily using digital calipers. All procedures adhered to the guidelines established by the York University Animal Care Committee (27). An infographic of this procedure can be seen in *Figure 1*.

**Figure 1:**
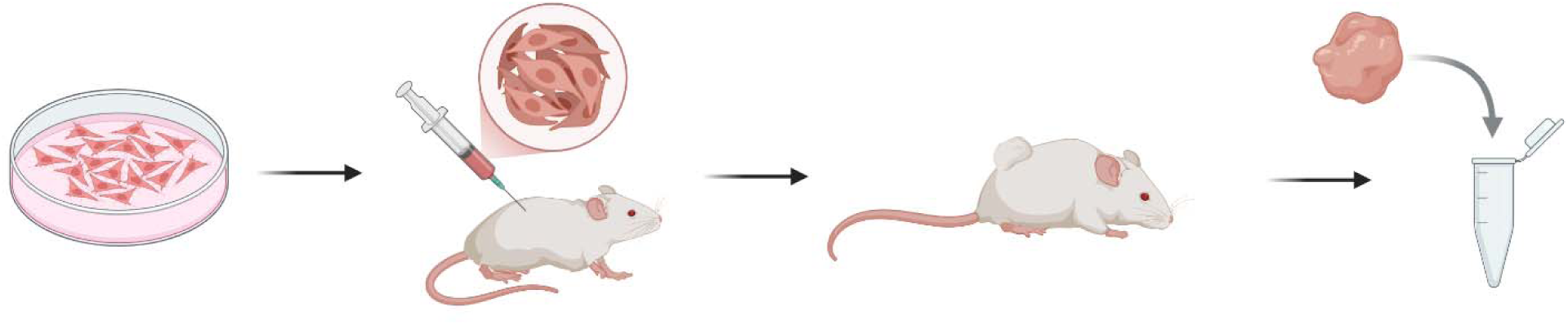
C26 mouse inoculation experimental design (Made with Biorender)

### Tissue Collection

At the endpoint of the study, and when mice were aged 10 to 12 weeks, the plantaris, soleus, red gastrocnemius, adipose tissue, liver and kidney were collected at week 4 under isoflurane anesthesia. Tumors were excised at both week 2 and week 4. All tissues were rapidly excised, weighed, and snap-frozen in liquid nitrogen and stored at –80°C for subsequent analysis [27].

### Tissue Homogenization

The skeletal muscle, liver and kidney tissue samples were weighed and homogenized in a 7-volume homogenization buffer. The tumor tissue samples were weighted and homogenized in a 5-volume homogenization buffer and the Adipose tissue samples were weighted and homogenized in a 1-volume homogenization buffer (composition: 20 mM HEPES, pH 7.4; 2 mM EGTA; 50 mM NaF; 100 mM KCl; 0.2 mM EDTA; 50 mM glycerophosphate), supplemented with 1 mM DTT, 1 mM benzamidine, 0.5 mM sodium vanadate, and protease and phosphatase inhibitor cocktails (10 μl/ml of buffer). The homogenization was performed on ice to maintain protein stability. The homogenate was centrifuged at 1,000 g for 3 minutes at 4°C, and the resulting supernatant was collected. For all tissues excluding Adipose, this supernatant was further centrifuged at 10,000 g for 30 minutes at 4°C to obtain clarified samples. The processed samples were subsequently used for western blotting and ultra-high-pressure liquid chromatography analyses.

### Protein Assay, Gels and Western Blotting of BCAA Metabolism

Protein concentrations were determined for each sample using the Pierce BCA Protein Assay Kit (Thermo Scientific, #23225). Absorbance readings were acquired at 550 nm using the KC4 plate reader software (Bio-Tek Instruments Inc., Winooski, Vermont, United States). A standard curve was generated using seven standards (0, 0.2, 0.4, 0.8, 1.0, 1.5, and 2.0 mg/mL) to calculate the sample volume required to load either 25 μg or 40 μg of protein per well, depending on the experimental conditions and sample amount.

Proteins were separated using 10% SDS-polyacrylamide gels with 10 or 15 wells per gel. Following electrophoresis, the separated proteins were transferred overnight onto 0.2 μm-pore polyvinylidene difluoride (PVDF) membranes (Bio-RAD, #1620177). Successful transfer was confirmed by staining membranes with Ponceau S dye for 15 minutes. Full Ponceau stained membranes were imaged using the BioRad ChemiDoc MP system. After imaging, membranes were followed by three 5-minute washes in Tris-buffered saline with Tween 20 (TBST).

To block non-specific antigen binding, membranes were incubated in a 5% milk solution (10 g skim milk powder in 190 mL TBST) at room temperature for 1 hour. After blocking, membranes were washed three times for 5 minutes each with TBST and incubated overnight at 4°C with the primary antibody of interest, prepared at the appropriate dilution in the blocking solution, as shown in *Table 1*.

**Table 1:**
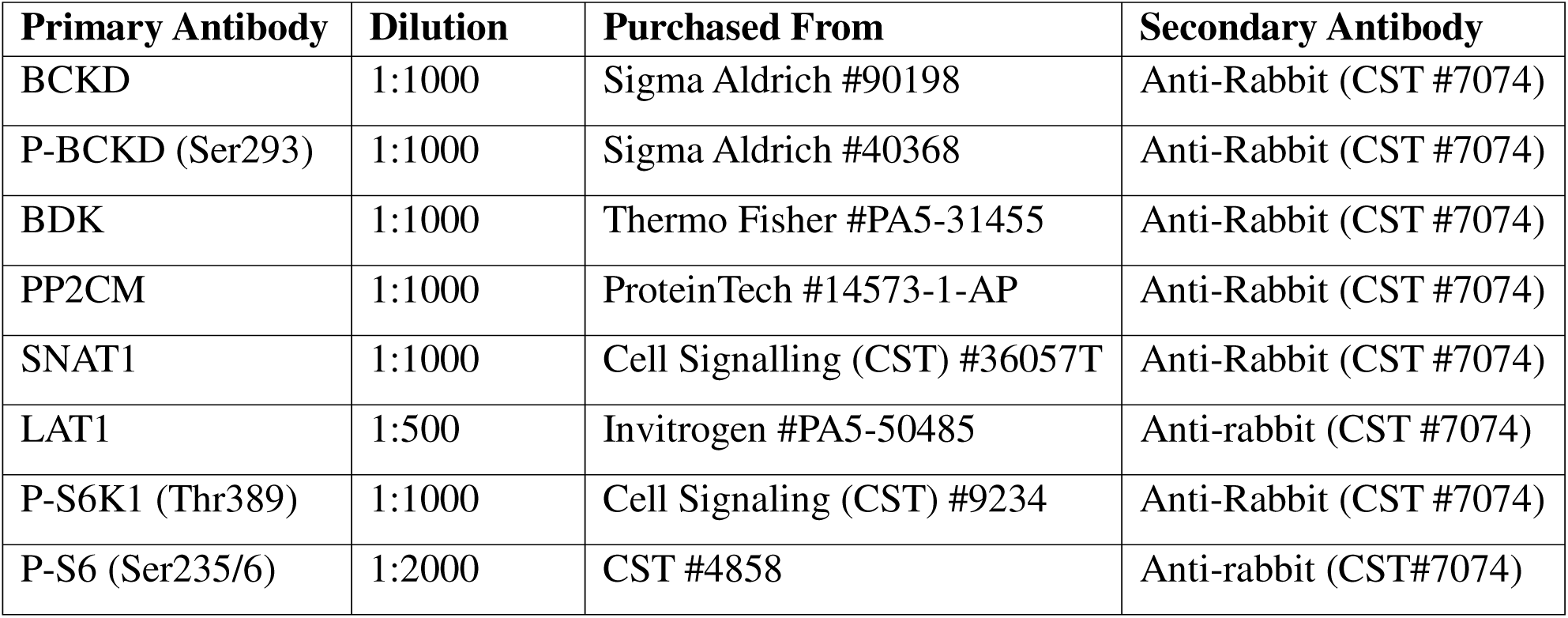
List of Antibodies Used.

The following day, membranes were washed again (3 × 5 minutes in TBST) and incubated with the appropriate secondary antibody (anti-rabbit, CST #7074) conjugated to HRP, and diluted at 1:10,000 in the blocking solution. Secondary antibody incubation was carried out for 3 hours at room temperature, after which membranes were washed again (3 × 5 minutes in TBST).

To visualize protein bands, HRP chemiluminescent substrate (Clarity Western ECL BioRad, #1705060S) was applied to each membrane. Signal detection and imaging were performed using the BioRad ChemiDoc MP system, and all images were quantified using Image Lab Software v7.

### BCKD assay

Frozen samples from red gastrocnemius, liver, kidney, adipose and tumor tissues were crushed in liquid nitrogen and homogenized using a Bio-Gen PRO200 Homogenizer. Homogenization was performed in 250μL of ice-cold buffer 1 (30 mM potassium phosphate, pH 7.4; 3 mM EDTA; 5 mM DTT; 1 mM valine; 3% FBS; 5% Triton X-100; and 1μM leupeptin). The homogenate was centrifuged at 10,000 g for 10 minutes at 4°C, and the supernatant was collected.

For the reaction, 50μL of the supernatant was added to 300μL of reaction buffer 2 (50 mM HEPES, pH 7.4; 30 mM potassium phosphate; 0.4 mM CoA; 3 mM NAD+; 5% FBS; 2 mM thiamine; 2 mM magnesium chloride; and 7.8μM [14C]-valine (Perkin Elmer, #NEC291EU050UC)). The reaction was carried out in a 1.5 mL Eppendorf tube with a 2 M NaOH soaked wick attached to the inner surface of the lid to capture released 14CO2. Tubes were tightly sealed with tape and incubated at 37°C for 30 minutes.

After incubation, the NaOH wick was removed and counted using a liquid scintillation counter to quantify the 14CO2 released during the BCKD reaction. Additionally, 50μL of the reaction mix was counted in the scintillation counter to measure total radioactivity. Protein concentrations were determined for each sample, and BCKD activity (measured from the wick) was normalized to total radioactivity and protein content.

Valine was used as the substrate for this assay, as valine-derived α-ketoisovalerate (KIV) is the preferred substrate for BCKD over leucine-derived intermediates.

### Analysis of Amino Acids by HPLC

#### Sample Preparation

Homogenized tissue samples containing free amino acids were diluted in a 1:2:1:8 ratio (sample: potassium phosphate buffer: 0.1 N hydrochloric acid: HPLC-grade water). The diluted samples were pre-column derivatized with a 1:1 ratio of sample to o-phthalaldehyde (OPA, 29.28 mM; Sigma Aldrich, #P1378) to enhance detection sensitivity.

Samples were injected into a YMC-Triart C18 column (C18, 1.9μm, 75 × 3.0 mm; YMC America, Allentown, PA, USA) fitted onto a Nexera X2 ultra-high-pressure liquid chromatography system (Shimadzu, Kyoto, Japan) equipped with a fluorescence detector. Detection was performed at an excitation wavelength of 340 nm and an emission wavelength of 455 nm.

Amino acids were separated using a gradient elution with two mobile phases: Mobile Phase A: 20 mM potassium phosphate buffer, pH 6.5; Mobile Phase B: 45% HPLC-grade acetonitrile, 40% HPLC-grade methanol, and 15% HPLC-grade water.

The gradient was programmed to increase from 5% to 100% Mobile Phase B over 21 minutes at a flow rate of 0.8 mL/min.

#### Quantification

Amino acid concentrations were calculated by integrating chromatographic peaks and comparing them to standard curves generated using BCAA and AA standards (Sigma Aldrich, #AAS18). For all samples, concentrations were normalized to total protein content.

### Analysis of Branched-Chain Ketoacids (BCKA) by HPLC

#### Sample Preparation

Tissue homogenates containing BCKAs were neutralized and diluted in a 1:2:1:8 ratio (sample: saturated potassium borate buffer: homogenization buffer or ddH O: ddH O). Standards were prepared using α-ketoisocaproate (KIC; Sigma Aldrich, #K0629), α-ketoisovalerate (KIV; Sigma Aldrich, #198994), and α-keto-β-methylvalerate (KMV; Sigma Aldrich, #K7125). Stock solutions (25 mg/mL) were serially diluted to generate standard curves, with HPLC-grade water serving as the blank. Samples and standards were filtered through 0.2 μm syringe filters (Sarstedt, #83.1826.001) prior to injection.

Samples were derivatized by adding 40 μL of 1,2-diamino-4,5-methylenedioxybenzene (DMB; Sigma Aldrich, #66807) solution to 40 μL of sample or standard, followed by incubation at 85 °C for 45 min in the dark. Reactions were cooled on ice for ≥5 min before analysis.

Derivatized samples were injected into an Intersil ODS-4 column (100 × 2.1 mm, 2 μm particle size; GL Sciences, #IN5020-81204) installed on a Nexera X2 UHPLC system (Shimadzu, Kyoto, Japan) equipped with a fluorescence detector. Detection was performed at excitation/emission wavelengths of 367/446 nm. Column temperature was maintained at 40 °C, with a sample compartment at 4 °C. Injection volume was 5 μL, and flow rate was 0.3 mL/min.

BCKAs were separated using a binary gradient of freshly prepared buffers: Mobile Phase A: 30% HPLC-grade methanol (Fisher Scientific, #A452-4), 70% HPLC-grade water; Mobile Phase B: 100% HPLC-grade methanol (Fisher Scientific, #A452-4).

The gradient program was as follows: 0–4 min, 0% B; 4–6 min, ramp to 50% B; 6–20 min, hold at 50% B. Maximum system pressure was maintained below 18,000 psi.

#### Quantification

BCKA concentrations (KIC, KIV, KMV) were calculated by integrating chromatographic peaks and comparing them to standard curves generated from external standards. Data were normalized to total protein concentration in each sample.

### Statistical Analysis

BCKD activity data were calculated as counts per minute (CPM) of CO released and bound to the wick, corrected for total radioactivity and normalized to the protein content (μg) of each sample. For western blot analyses, protein loading was normalized using Ponceau values.

Statistical analyses were performed using GraphPad Prism software (versions 7 or 8; GraphPad, Massachusetts, United States). Data are expressed as mean ± standard error of the mean (SEM). Unpaired t-tests were used to compare group means. Statistical significance was set at p < 0.05.

## Results

To confirm that mice had entered a cachectic, muscle-wasting state, parameters such as body weight, tumor-free body weight, and skeletal muscle mass were evaluated during the original animal experiments and confirmed significant reductions in the weights of those organs and tissues (27).).

### Tumor

Previous studies have reported that tumors may exhibit increased uptake and accumulation of BCAAs to support their rapid growth and proliferation (22,23). Given these findings, we sought to determine whether similar changes occur in the C26 tumor model. To do this, we assessed BCAA, BCKA and total AA content using HPLC. We also examined BCKD enzymatic activity capacity and used western blotting to evaluate protein expression of key components of BCAA metabolism, including BCKD regulatory enzymes, AA transporters (LAT1 and SNAT1), and the mTORC1 signaling marker P-S6. This multi-level analysis provides insight into how the tumor alters AA handling to support its metabolic demands.

#### Tumor tissue shows increased BCAA levels and BCKD activity at Week 4

HPLC analysis revealed significantly elevated concentrations of valine (p < 0.001), isoleucine (p < 0.05), and total BCAAs (p < 0.05) in Week 4 tumors compared to Week 2 (**Figure 2A**), while leucine levels showed a trend toward an increase (p = 0.09). BCKA levels were unchanged between the groups (Fig 2**B**). Additionally, while some AAs, such as histidine, arginine, phenylalanine, and proline appeared modestly elevated at Week 4, none reached statistical significance (Fig 2**C**). Glutamate was also increased by 65% at Week 4, though this was not statistically significant (Fig 2**D**).

**Figure 2:**
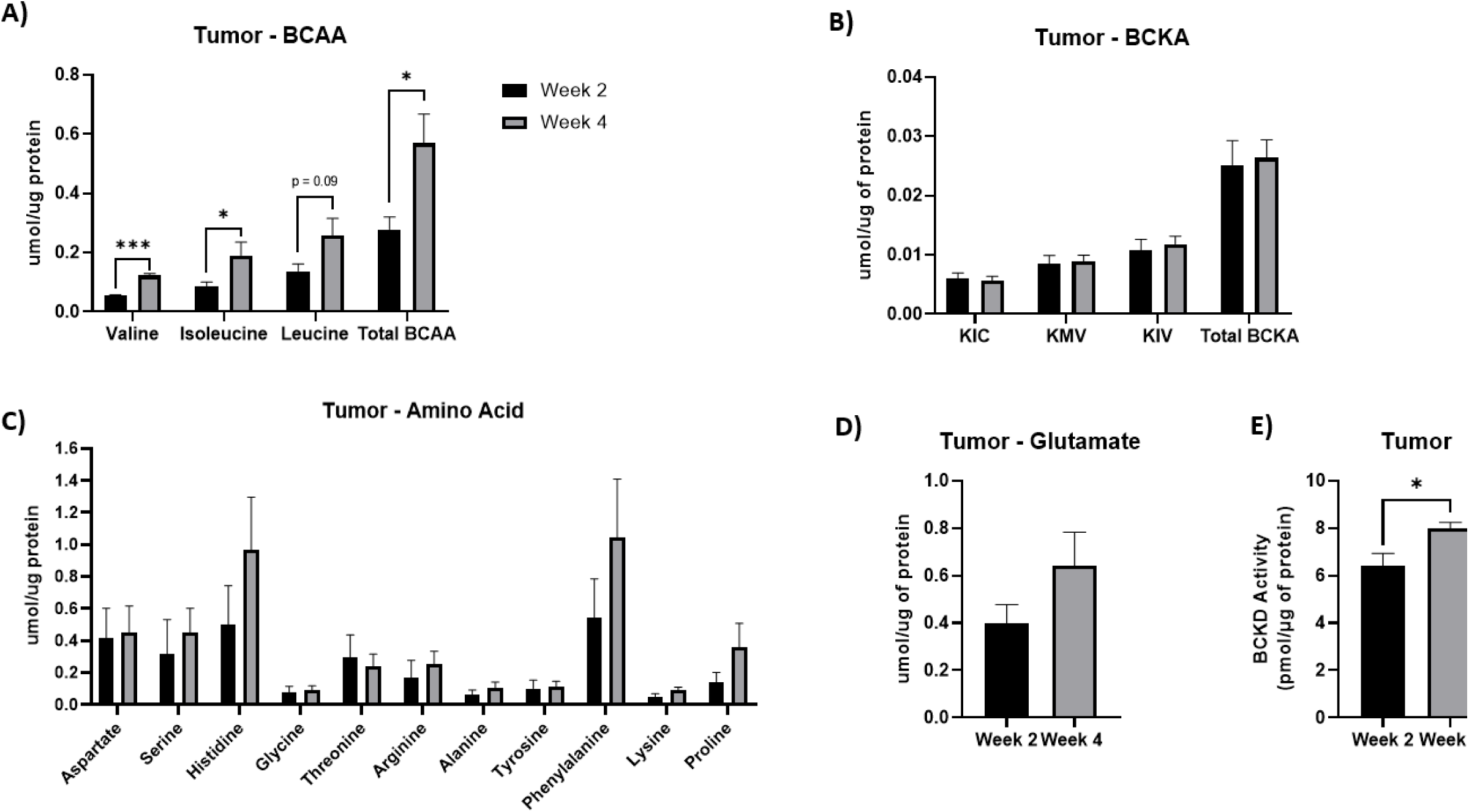
Increased BCAA concentrations and BCKD activity in C26 tumor tissue at week 4. Valine, isoleucine, leucine, total BCAAs (**A**), kic, kmv, kiv and total BCKAs (**B**), other AAs (**C),** glutamate (**D**) and BCKD activity assay (**E**) were measured in tumor tissue samples collected at Week 2 (n = 5) and Week 4 (n = 10) following C26 cell inoculation. AA and BCKA concentrations (**A-D**) are expressed in µmol per µg of protein. BCKD activity (**E**), measured using [^14^C]-CO_2_ released per µg of protein per minute. Statistical significance was determined using unpaired t-tests. Data are presented as mean ± SEM. ***p < 0.001, *p < 0.05.

Further, to assess whether increased BCAA levels were accompanied by altered catabolic activity, we measured the activity of the BCKD complex. BCKD activity was significantly elevated (p < 0.05) in Week 4 tumors (Fig 2**E**), suggesting that in addition to accumulating BCAAs, the tumor may also upregulate their oxidation to support its metabolic demands.

#### Upregulation of PP2CM, SNAT1 and P-S6 in tumor-tissue at week 4

To explore the mechanism underlying increased AA concentration and BCKD activity in tumor tissue, we assessed the expression of key BCAA catabolic enzymes, transporters, and signaling proteins by western blotting (**Figure 3A**). While there was no statistically significant difference in BCKD abundance between tumors collected at Week 2 and Week 4 (Fig 3**B**), PP2CM, the phosphatase responsible for BCKD activation, was significantly upregulated (p < 0.05) at Week 4 (Fig 3**C**). In contrast, P-BCKD (Fig 3**D**) and BDK (Fig 3**E**) protein levels were not significantly different between groups. These findings suggest that the observed increase in BCKD activity is likely due to the increase in PP2CM. LAT1 expression was not significantly different between timepoints (Fig 3**F**), whereas SNAT1 was significantly upregulated (p < 0.01) at Week 4 (Fig 3**G**), indicating increased transporter expression specific to certain AA pathways. Expression of P-S6, a downstream effector of mTORC1 signaling, was also significantly elevated (p < 0.05) at Week 4 (Fig 3**H**), suggesting enhanced nutrient-driven signaling and protein synthesis during tumor progression. Collectively, these results indicate that tumor growth is accompanied by selective upregulation of BCAA catabolic regulation, AA transport, and mTORC1 activation.

**Figure 3:**
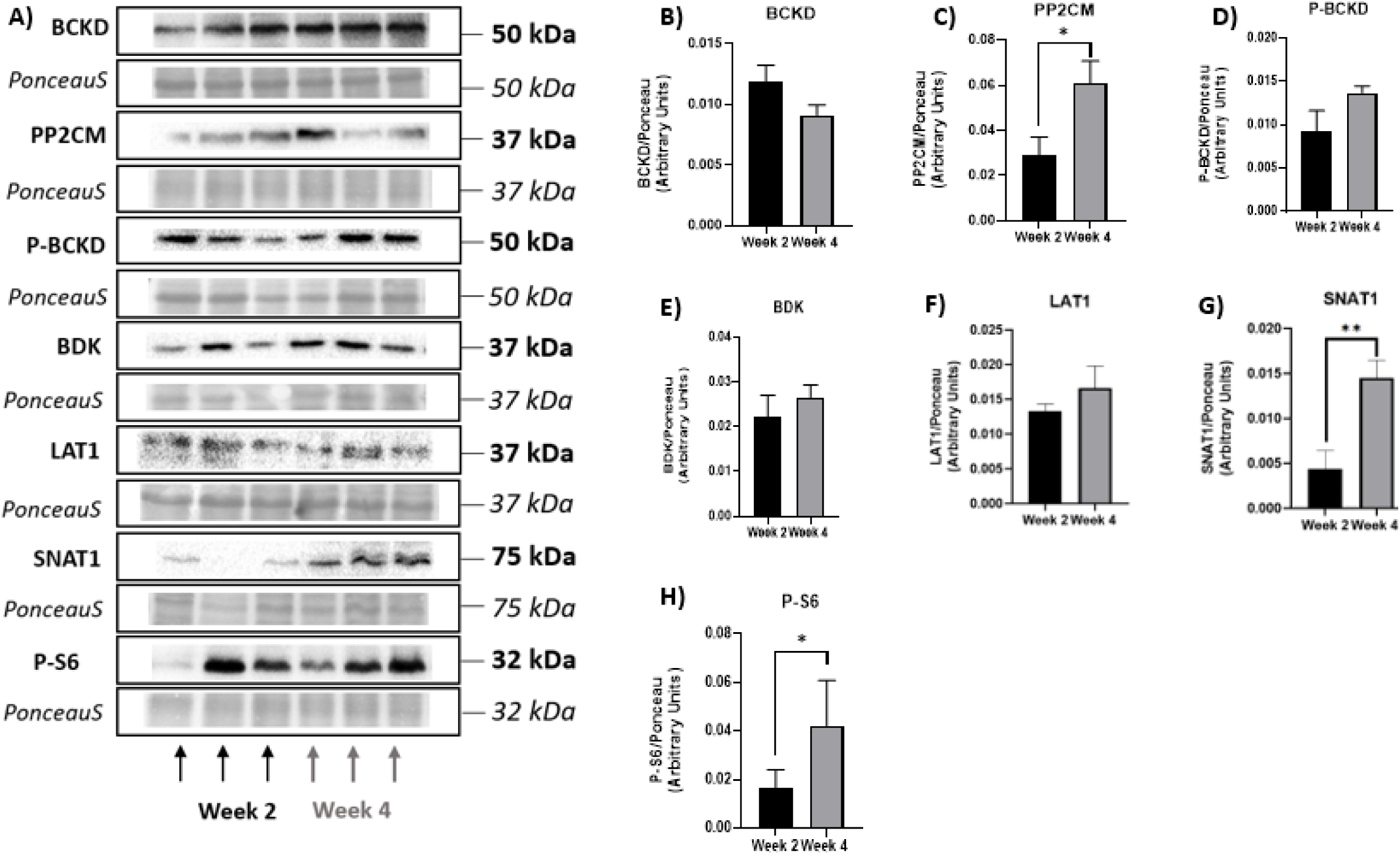
Upregulation of PP2CM, SNAT1, and P-S6 expression in tumor tissue at week 4. Representative blots (**A**) and quantified protein expression of BCKD (**B**), PP2CM (**C**), P-BCKD (**D**), BDK (**E**), LAT1 (**F**), SNAT1 (**G**), and P-S6 (**H**) were measured by western blotting in tumor tissue collected at Week 2 (n = 5) and Week 4 (n = 10). Protein levels were normalized to Ponceau staining. Data are presented as mean ± SEM. Statistical significance was determined using unpaired t-tests. **p < 0.01, *p < 0.05.

### Liver

Given the significant increase in BCAA content, enzyme activity, SNAT1 and P-S6 expression, observed in the week 4 tumors, we next examined the liver, a central organ in AA metabolism and homeostasis (30). The liver plays a key role in BCAA catabolism via the BCKD complex (31,32) and has been implicated in systemic metabolic adaptations during cancer progression (33).

#### Altered AA concentrations in liver of C26 tumor-bearing mice

Liver BCAAs (Figure 4**A**), BCKAs (Fig 4**B**) and glutamate (Fig 4**D**) concentrations were not significantly different between groups, suggesting preserved hepatic BCAA content during tumor progression. Several other AAs were altered as aspartate (p < 0.001), arginine (p < 0.001), and alanine (p < 0.01) were significantly decreased, while lysine was significantly increased (p < 0.01) (Fig 4**C**). These changes suggest that tumor burden may contribute to selective hepatic AA depletion, particularly affecting NEAAs. BCKD activity was decreased by 20% in the C26 group, however, was not statistically significant (Fig 4**E**), indicating a modest suppression of hepatic BCAA catabolism in response to tumor burden.

**Figure 4:**
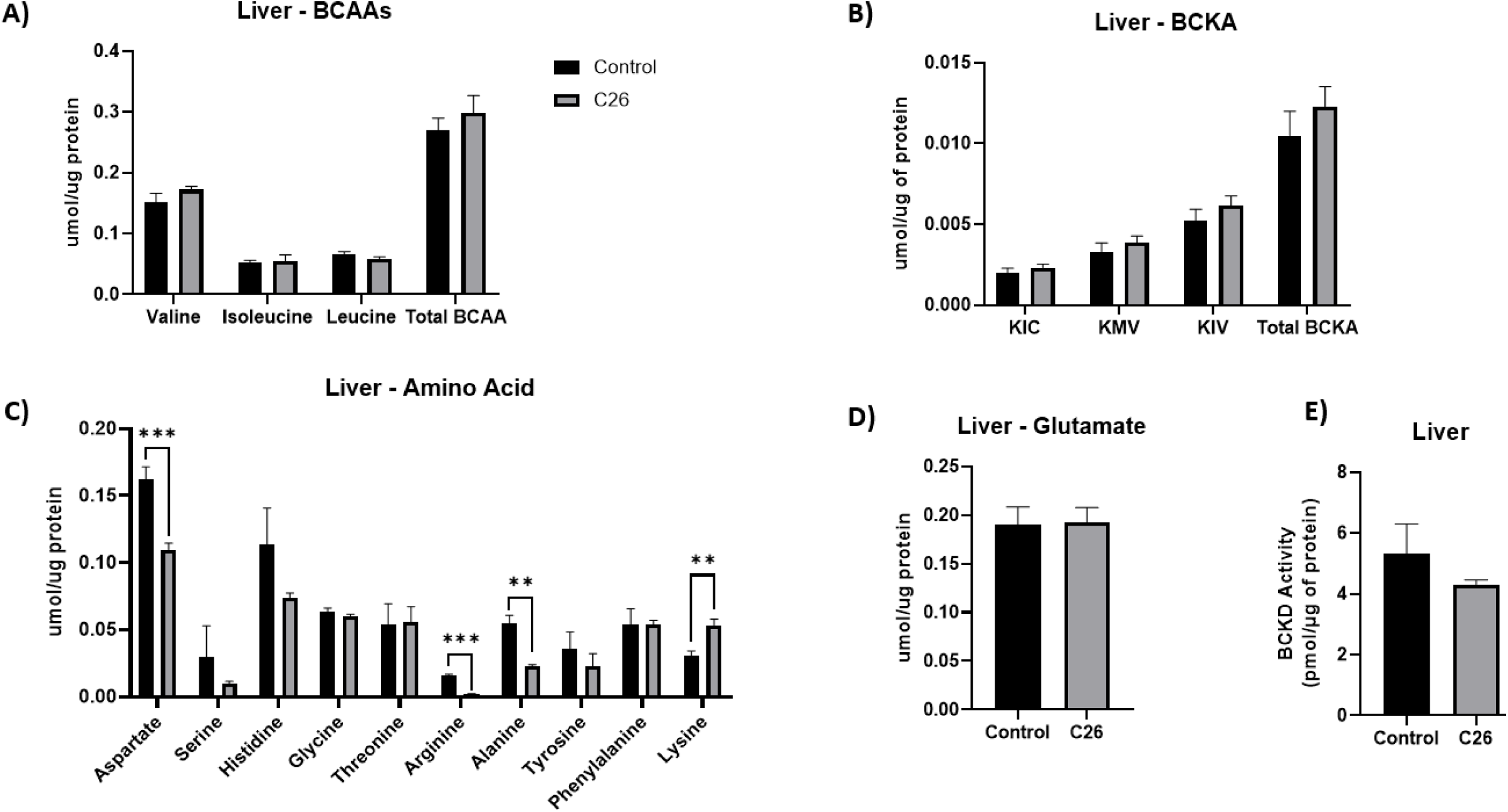
Altered AA concentrations in liver tissue of C26 tumor-bearing mice. BCAA concentrations (**A**), BCKA concentrations (**B**), other AAs (**C**), glutamate (**D**), and BCKD activity (**E**) were measured in liver tissue collected from control (n = 10) and C26 tumor-bearing (n = 10) mice. AA nd BCKA (**A–D**) are expressed in µmol per µg of protein. BCKD activity (**E**), measured using [¹ C]-valin, is expressed as pmol of [¹ C]-CO released per µg of protein per minute. Data are presented as mean ± SEM. Statistical significance was determined using unpaired t-tests. ****p < 0.001, **p < 0.01, *p < 0.05.

Upregulated PP2CM, P-BCKD, LAT1 and P-S6 expression in the liver of C26 tumor-bearing mice.

BCKD expression was not significantly different between groups (**Figure 5A-B**), while PP2CM was significantly elevated (p < 0.01) in the C26 group (Fig 7A, **C**). P-BCKD showed a trend toward increased expression (p = 0.06) (Fig 5A, **D**), and BDK levels were unchanged (Fig 5A, **E**), suggesting a shift in hepatic BCAA regulatory balance during tumor progression. LAT1 expression was significantly decreased (p < 0.05) in tumor-bearing mice (Fig 5A, **F**), while SNAT1 was unchanged (Fig 5A, **G**). P-S6 expression trended higher (p = 0.06) in the C26 group (Fig 5A, **H**), indicating potential upregulation of hepatic mTORC1 signaling.

**Figure 5:**
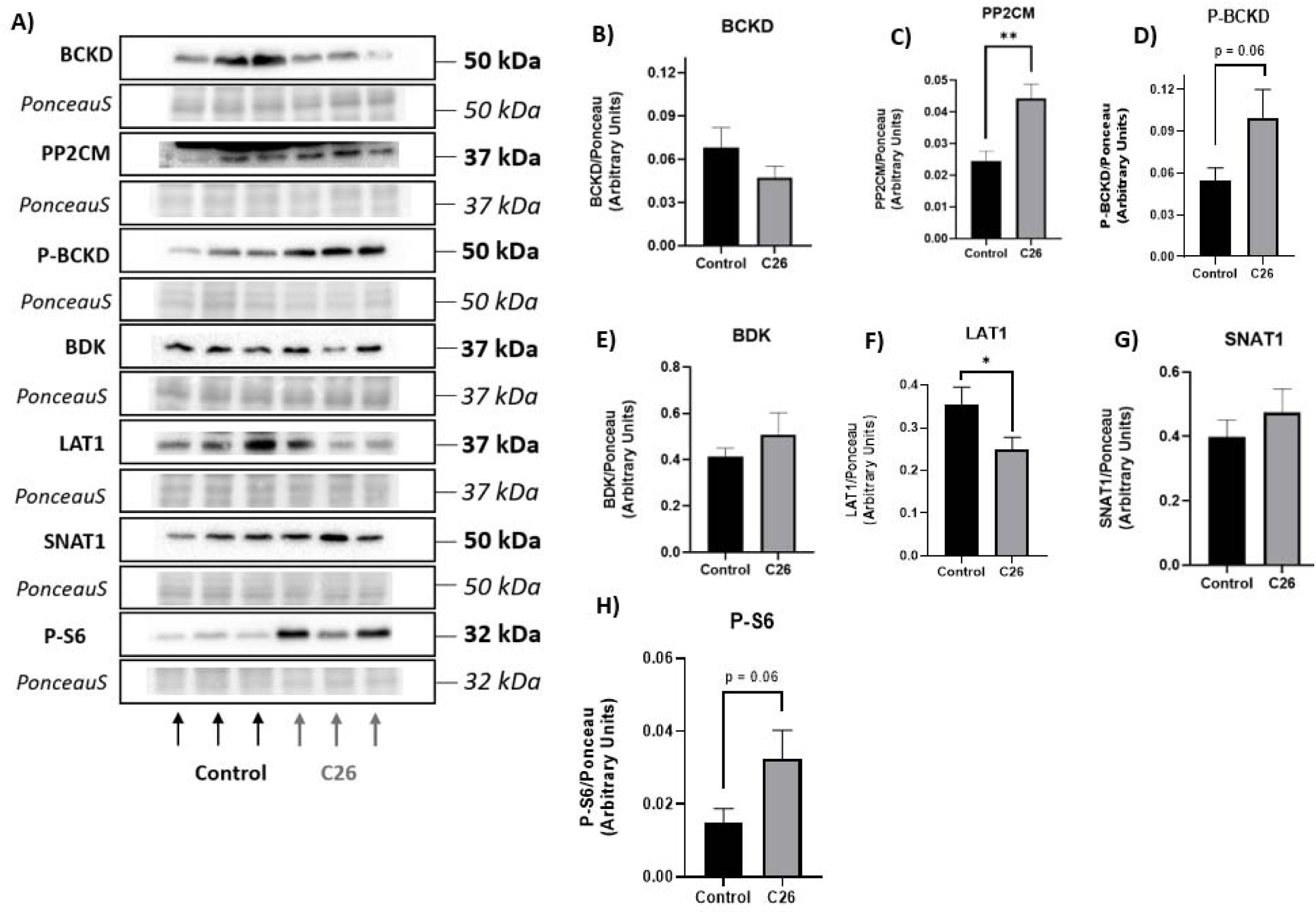
Upregulated PP2CM, P-BCKD, LAT1 and P-S6 expression in the liver tissue of C26 tumour-bearing mice. Representative blots (**A**) and quantified protein expression of BCKD (**B**), PP2CM (**C**), P-BCKD (**D**), BDK (**E**), LAT1 (**F**), SNAT1 (**G**), and P-S6 (**H**) were measured by western blotting in liver tissue collected from control (n = 10) and C26 tumor-bearing (n = 10) mice. Protein levels were normalized to Ponceau staining. Data are pr sented as mean ± SEM. Statistical significance was determined using unpaired t-tests. **p < 0.01, *p < 0.05.

### Kidney

The kidney plays an important yet underappreciated role in systemic AA metabolism, including BCAA catabolism, ammonia production, and gluconeogenesis (34). Although the kidney is not typically central in cachexia research, its involvement in maintaining nitrogen balance, and AA homeostasis (35) suggests it may contribute to the systemic metabolic disturbances observed in cachexia. Emerging evidence also indicates that kidney metabolism may become metabolically altered in disease states such as cancer (36), warranting further exploration of its role in cancer-associated muscle wasting.

#### Decreased valine, BCKAs and alanine concentration is kidney tissue of C26 tumor-bearing mice

HPLC analysis showed a significant reduction in valine levels (p < 0.01) in the C26 group, while isoleucine, leucine, and total BCAAs were unchanged (Figure 6**A**). BCKAs were also reduced, with significant decreases in KIC, KMV, and KIV (p < 0.05), and total BCKAs (p = 0.06) in the C26 group (Fig 6**B**). Among other AAs, arginine showed a trend toward reduction (p = 0.06), and alanine was significantly decreased (p < 0.05), while no other AAs differed between groups (Fig 6**C**). Glutamate concentrations were unchanged (Fig 6**D**). Finally, BCKD activity was similar between groups (Fig 6**E**), indicating stable renal BCAA catabolism during tumor progression.

**Figure 6:**
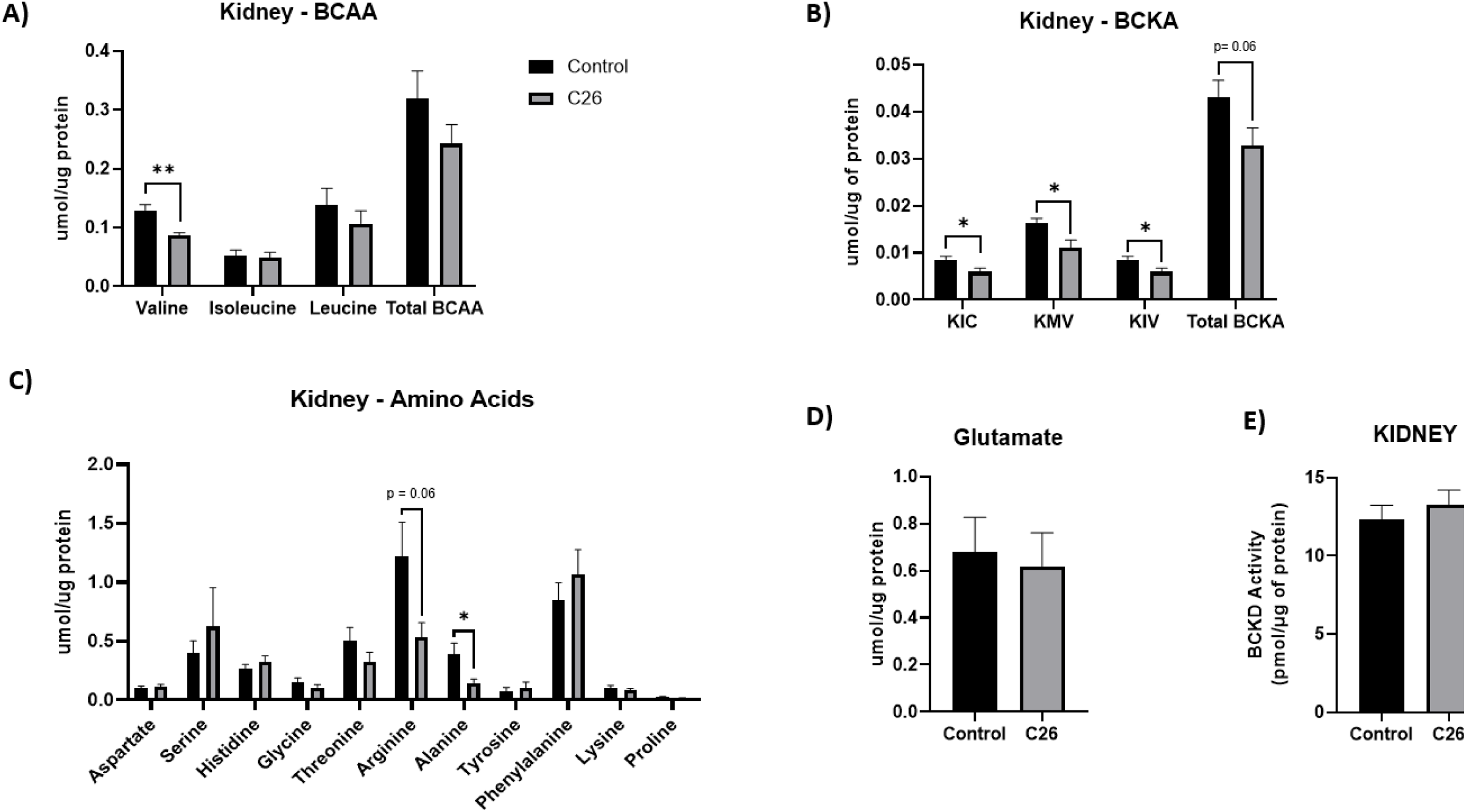
Decreased valine, BCKAs and AA concentrations in kidney tissue of C26 tumor bearing mice. BCAA concentrations (**A**), BCKA concentrations (**B**), other AAs (**C**), glutamate (D), and BCKD activity (**E**) were measured in kidney tissue collected from control (n = 10) and C26 tumor-bearing (n = 9) mice. AA and BCKA concentrations (**A–D**) are expressed in µmol per µg of protein. BCKD activity (**E**), measured using [¹ C]-valine, is expressed as pmol of [¹ C]-CO released per µg of protein per minute. Data are presented as mean ± SEM. Statistical significance was determined using unpaired t-tests. **p < 0.01, *p < 0.05.

#### Tumor burden alters expression of BCKD regulators and AA transporters in kidney tissue

Western blotting analysis revealed that while BCKD (**Figure 7A-B**) and BDK (Fig 7A, **E**), levels remained unchanged. significant decreases in both PP2CM (p < 0.001) (Fig 7A, **C**) and P-BCKD (p < 0.01) (Fig 7A, **D**) were observed. These changes may reflect a regulatory balance that allows BCKD activity within the kidney to remain stable, as observed in **Figure 6E**. Furthermore, LAT1 expression was significantly decreased (p < 0.01) (**Fig 7A, F**), whereas SNAT1 was significantly upregulated (0.001) (Fig 7**G**), suggesting altered AA transport. P-S6 expression was unchanged (Fig 7A, **H**), indicating no major shifts in mTORC1 signaling at this time point.

**Figure 7:**
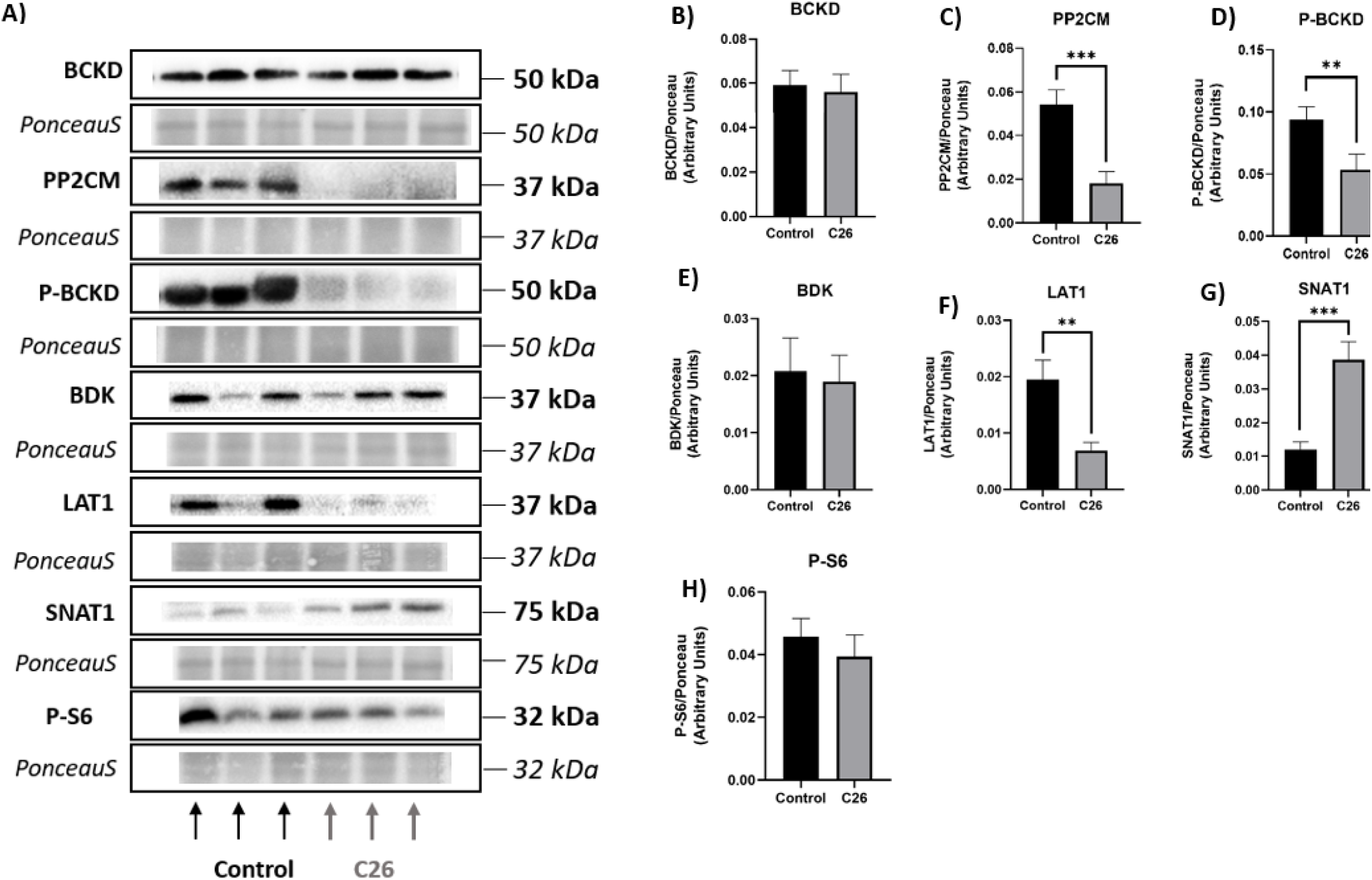
Decreased PP2CM, P-BCKD, LAT1 and increased SNAT1 expression in kidney tissue of C26 tumor-bearing mice. Representative blots (**A**) and quantified protein expression of BCKD (**B**), PP2C (**C**), P-BCKD (**D**), BDK (**E**), LAT1 (**F**), SNAT1 (**G**), and P-S6 (**H**) were measured by western blotting in kidney tissue collected from control (n = 10) and C26 tumor-bearing (n = 9) mice. Protein levels were normali ed to Ponceau staining. Data are presented as mean ± SEM. Statistical significance was determined using un aired t-tests. ***p < 0.001, *p < 0.05.

### Adipose

Adipose tissue, beyond its role in energy storage, contributes to whole-body AA metabolism and inter-organ communication (37). It also plays a role in cancer and cachexia, where inflammation and lipolysis disrupt metabolic homeostasis and contribute to systemic wasting (38). Given the changes observed in tumor, liver, and kidney, we next assessed adipose tissue.

#### Decreased glutamate and altered AA concentrations in adipose tissues of C26 tumor-bearing mice

BCAA concentrations were unchanged between groups (**Figure 8A**), Adipose BCKAs were also not significantly different between control and C26 mice (Fig 8**B**). Several other AAs were altered, with significant reductions in aspartate and glycine (p < 0.001), threonine (p < 0.01), alanine, and lysine (p < 0.001), while tyrosine was significantly increased (p < 0.001) and serine showed a trend toward elevation (p = 0.07) (Fig 8**C**). Glutamate levels were significantly reduced in C26 adipose tissue (p < 0.001) (Fig 8**D**), suggesting altered nitrogen handling or transamination activity in response to tumor burden. BCKD activity was unchanged (Fig 8**E**), suggesting that adipose BCAA catabolism remains relatively stable despite these molecular changes.

**Figure 8:** Decreased glutamate and altered AA concentrations in adipose tissues of C26 tumor-bearing mice. BCAA concentrations (**A**), BCKA concentrations (**B**), other AAs (**C**), glutamate (**D**), and BCKD activity (**E**) were measured in adipose tissue collected from control (n = 10) and C26 tumor-bearing (n = 10) mice. AA BCKA concentrations (**A–D**) are expressed in µmol per µg of protein. BCKD activity (**E**), measured using [¹ C]-valine, is expressed as pmol of [¹ C]-CO released per µg of protein per minute. Data are presented as mean ± SEM. Statistical significance was determined using unpaired t-tests. ***p < 0.001, **p < 0.01, *p < 0.05.

#### Reduced LAT1 and P-S6 expression in adipose tissue of C26 tumor-bearing mice

Expression of BCKD, PP2CM, and P-BCKD was not significantly different between groups (**Figure 9A–D**), while BDK expression was elevated in the C26 group, approaching statistical significance (p = 0.06) (Fig 9A, **E**). This increase in BDK may reflect a subtle regulatory shift, though P-BCKD remained unchanged (Fig 9A**, D**), suggesting that adipose BCAA oxidation is maintained despite altered enzyme expression. LAT1 (Fig 9A, **F**) and P-S6 (Fig 9A, **H**) expression were significantly reduced (p < 0.01) in the C26 group, while SNAT1 levels decreased by 30%, however, was not statistically significant (Fig 9A, **G**). These changes suggest that tumor burden is associated with decreased AA transport and suppressed mTORC1 signaling in adipose tissue.

**Figure 9:**
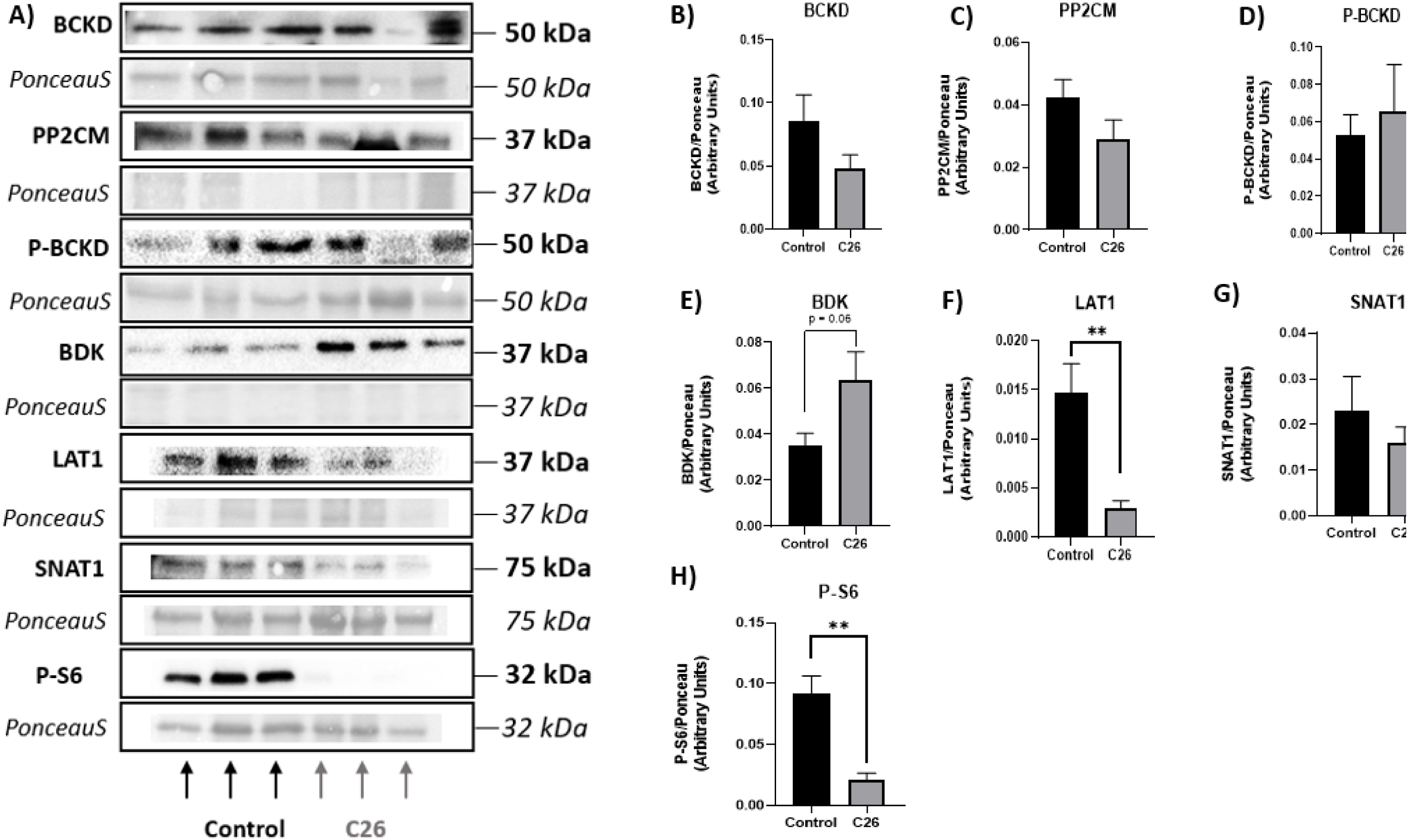
Decreased LAT1 and P-S6 expression in adipose tissue of C26 tumor-bearing mice. Representative blots (**A**) and quantified protein expression of BCKD (**B**), PP2CM (**C**), P-BCKD (**D**), BDK (**E**), LAT1 (**F**), SNAT1 (**G**), and P-S6 (**H**) were measured by western blotting in adipose tissue collected from control (n = 10) and C26 tumor-bearing (n = 10) mice. Protein levels were normalized to Ponceau staining. Data are presented as mean ± SEM. Statistical significance was determined using unpaired t-tests. **p < 0.01.

### Skeletal Muscle

Skeletal muscle is a major site of BCAA metabolism and a key tissue affected by cancer cachexia (39, 40). Given its central role in whole-body protein turnover and energy expenditure, skeletal muscle is a critical target for understanding how tumor burden impacts systemic AA handling and nutrient signaling. Cachexia-associated muscle loss is particularly detrimental to quality of life and survival (8), making it essential to examine tissue-specific metabolic adaptations in different muscle types.

To investigate these effects, three skeletal muscles with varying fiber-type composition and metabolic profiles were selected: red gastrocnemius, soleus, and plantaris. These muscles offer insights into fiber-type-specific responses to tumor burden and their potential contributions to whole-body metabolic remodeling in cancer.

### Red Gastrocnemius

The red portion of the gastrocnemius muscle is composed predominantly of oxidative, slow type I and fast type IIa fibers, which are rich in mitochondria and heavily involved in aerobic metabolism (41,42). This fiber composition makes red gastrocnemius particularly relevant for examining metabolic shifts during tumor progression, due to its high mitochondrial content, role in BCAA metabolism and BCAA sensitive mTORC1 signalling (43,44). Together, these features make it a useful indicator of cancer-induced disruptions in nutrient and energy metabolism.

#### Reduced AA concentrations and BCKD activity in red gastrocnemius of tumor-bearing mice

HPLC analysis revealed significantly lower concentrations of valine, isoleucine, leucine, and total BCAAs (p < 0.05) in the red gastrocnemius of C26 tumor-bearing mice (Figure 10**A**). BCKA concentrations, however, were unchanged between groups (Fig 10**B**). Among other AAs, alanine (p < 0.01) and lysine (p < 0.001) were significantly decreased, while no other AAs showed significant differences (Fig 10**C**). Glutamate levels were unchanged (Fig 10**D**). Finally, BCKD activity was significantly reduced in the C26 group (p < 0.01) (Fig 10**E**), suggesting that the observed decline in BCAAs is not due to increased local catabolism, however, may reflect systemic redistribution, towards tumor tissue. The red gastrocnemius may also be attempting to preserve its limited BCAA pool by downregulating BCKD activity.

**Figure 10:**
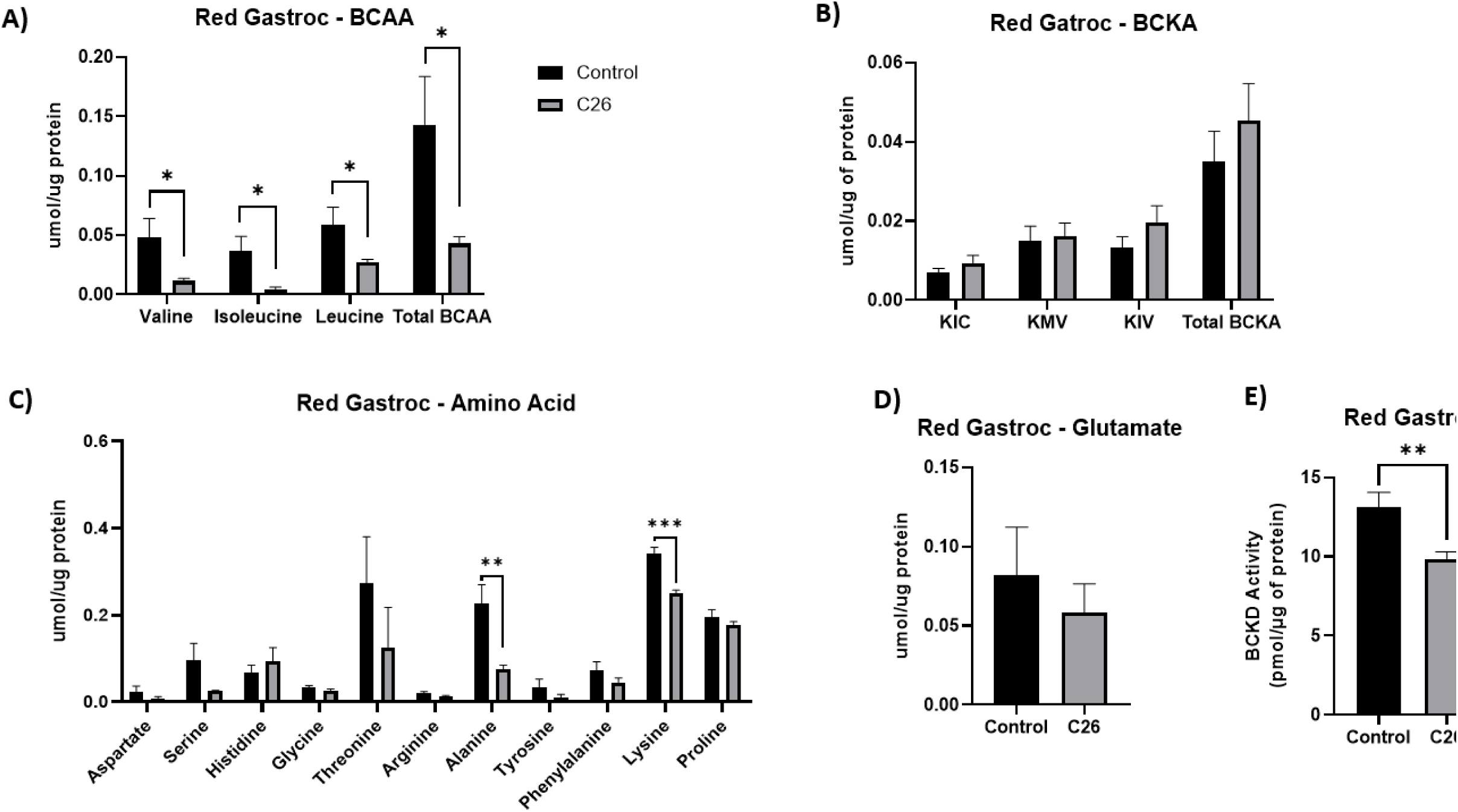
Decreased BCAA, AA concentrations and BCKD activity in red gastrocnemius muscle of C26 tumor-bearing mice. BCAA concentrations (**A**), BCKD concentrations (**B**), other AAs (**C**), glutamate (**D**), and BCKD activity (**E**) were measured in red gastrocnemius muscle tissue collected from control (n = 10) and C26 tumor-bearing (n = 10) mice. AA and BCKD concentrations (**A–D**) are expressed in µmol per µg of protein. BCKD activity (**D**), measured using [¹ C]-valine, is expressed as pmol of [¹ C]-CO released per µg of protein per minute. Data are presented as mean ± SEM. Statistical significance was determined using unpaired t-tests. ***p < 0.001, **p < 0.01, *p < 0.05.

#### Decreased LAT1 and P-S6 expression in red gastrocnemius muscle of C26 tumor-bearing

Western blotting revealed no significant differences in the expression of BCKD, PP2CM, P-BCKD, or BDK between groups (**Figure 11A-E**), suggesting that the reduction in BCKD activity observed in **Figure 10E** is not due to altered protein abundance or phosphorylation, however, may be due to the lower levels of BCAAs resulting in less activation of the BCKD complex. LAT1 expression was significantly reduced (p < 0.001) in the C26 group (Fig 11A, **F**), while SNAT1 remained unchanged (Fig 11A, **G**). P-S6 expression was also significantly lower (p < 0.05) in tumor-bearing mice (Fig 11A, **H**), indicating suppression of mTORC1 signaling. Together, these changes suggest that reduced BCAA uptake and suppressed anabolic signaling may contribute to skeletal muscle wasting during tumor progression.

**Figure 11:**
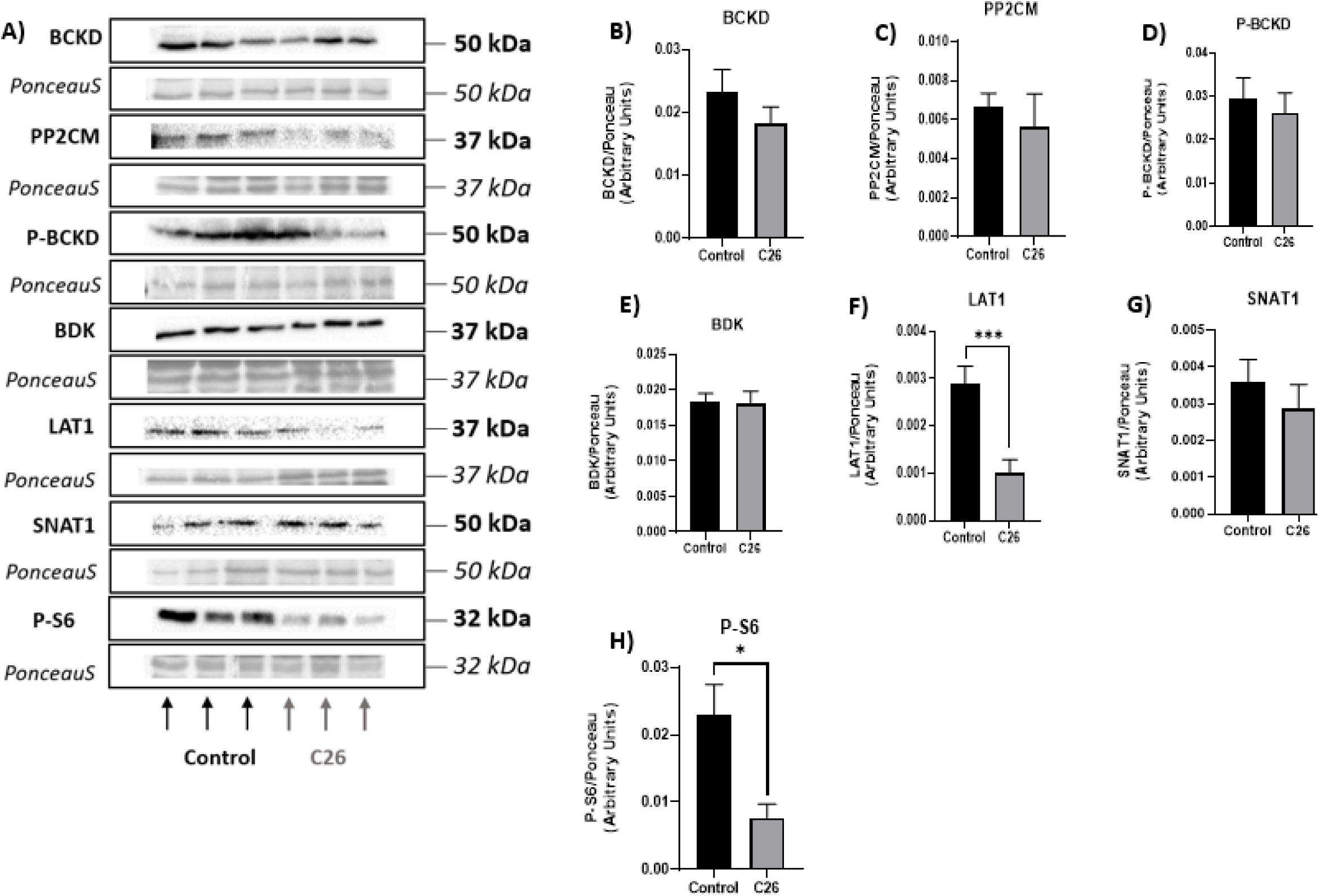
Decreased LAT1 and P-S6 expression in red gastrocnemius muscle of C26 tumor-bearing mice. Representative blots (**A**) and quantified protein expression of BCKD (**B**), PP2CM (**C**), P-BCKD (**D**), BDK (**E**), LAT1 (**F**), SNAT1 (**G**), and P-S6 (**H**) were measured by western blotting in red gastrocnemius muscle tissue collected from control (n = 10) and C26 tumor-bearing (n = 10) mice. Protein levels were normalized to Ponceau staining. Data are presented as mean ± SEM. Statistical significance was determined using unpaired t-tests. ***p < 0.001, *p < 0.05.

### Plantaris

Following the findings in red gastrocnemius, we next examined the plantaris muscle, which is located in the lower hindlimb and plays a role in locomotion (45). The plantaris muscle is predominantly composed of fast-twitch fibers, including type IIB, IIX/IIB hybrids, and IIX fibers, with a smaller proportion of oxidative type IIA fibers (41,42). This contrasts with muscles like the soleus, which are enriched in slow-twitch fibers (Type I and IIA) and are more fatigue-resistant (41,43). The plantaris is therefore considered a fast muscle with a high glycolytic capacity and moderate oxidative potential (41,42). Its mixed fiber-type composition makes it an informative tissue for evaluating intermediate responses to tumor-induced metabolic stress. By examining BCAA metabolism and related pathways in the plantaris, we aimed to determine whether the changes observed in red gastrocnemius were muscle-specific or reflected broader trends across skeletal muscle during cancer progression.

#### Decreased BCAA and proline concentrations in plantaris muscle of tumor-bearing mice

HPLC analysis revealed significantly lower levels of isoleucine (p < 0.01), leucine (p < 0.05), and total BCAAs (p < 0.05) in plantaris muscle of C26 tumor-bearing mice (**Figure 12A**). Although glutamate levels were not significantly different, there was a 157% increase in the C26 group (Fig 12**B**). Among other AAs proline was significantly decreased (p < 0.05), while threonine showed a non-significant reduction of 49%. Glycine and phenylalanine showed non-significant elevation of 173% and 166% respectively (Fig 12**C**). These findings suggest that BCAA depletion extends beyond red gastrocnemius and is accompanied by selective changes in other AAs in the plantaris during tumor progression.

**Figure 12:**
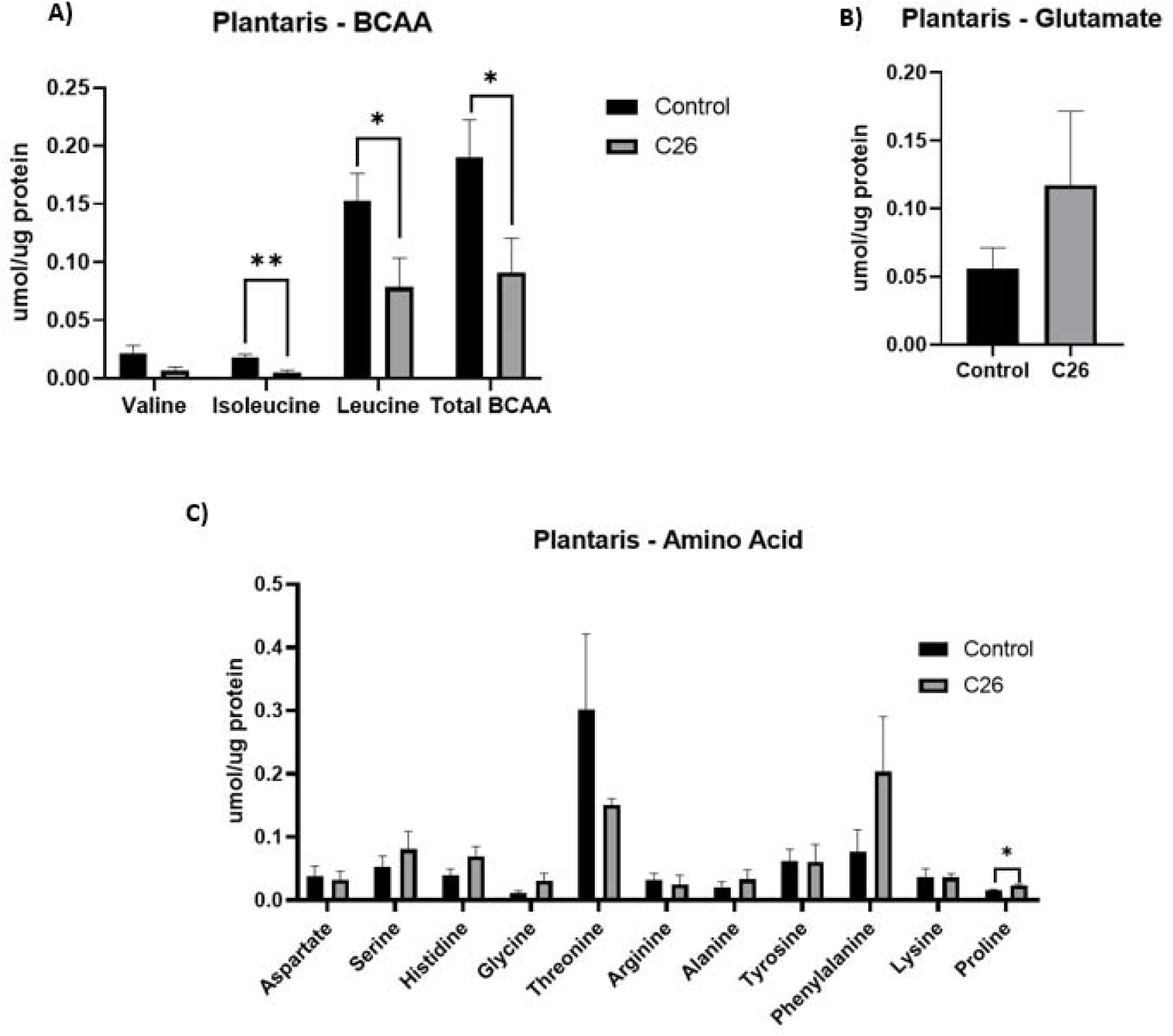
Decreased BCAA and proline concentrations in plantaris muscle of C26 tumor-bearin mice. BCAA concentrations (**A**), glutamate (**B**), and other AAs (**C**) were measured in plantaris muscle tissue collected from control (n = 10) and C26 tumor-bearing (n = 10) mice. AA concentrations are expressed in µmol per µg of protein. Data are presented as mean ± SEM. Statistical significance was determined using unpaired t-tests. **p < 0.01, *p < 0.05.

#### Trends toward reduced LAT1 and P-S6K1 expression in plantaris muscle of tumor-bearing mice

No significant differences in the expression of BCKD, PP2CM, P-BCKD, or BDK between groups were found (Fig 13**–E**), suggesting that the observed reduction in BCAAs in **Figure 12A** is not due to altered expression of BCAA metabolic enzymes. LAT1 (Fig 13A, F) and P-S6K1 (Fig 13A, H) expression were lower in the C26 group, however, did not reach statistical significance (p = 0.09 and p = 0.08, respectively, while SNAT1 remained unchanged (Fig 13A, **G**). These trends are consistent with changes observed in red gastrocnemius and may reflect early suppression of AA transport and anabolic signaling in plantaris muscle during tumor progression.

**Figure 13:**
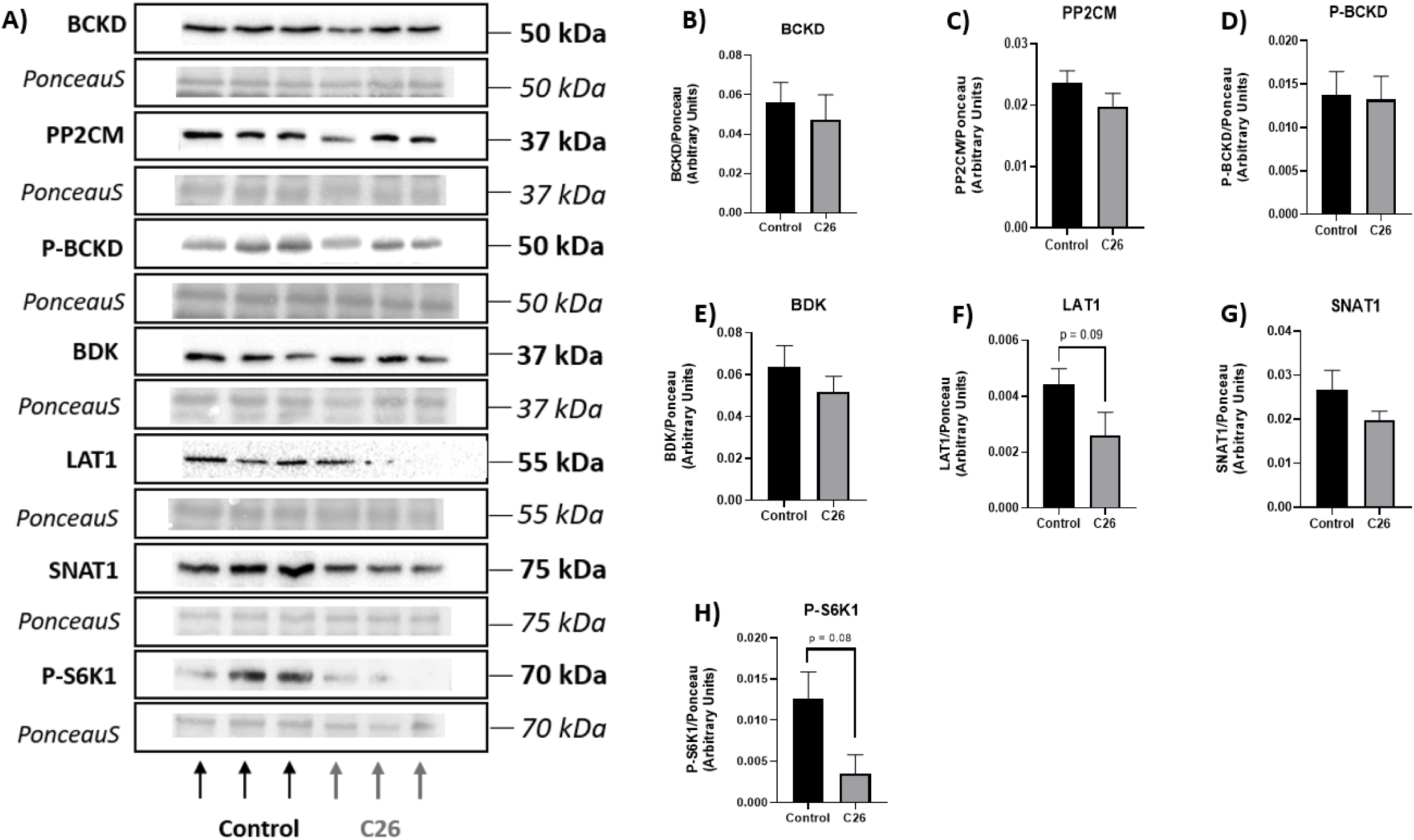
Trends for decreased LAT1 and P-S6K1 expression in the plantaris muscle of C26 tumo - bearing mice. Representative blots (**A**) and quantified protein expression of BCKD (**B**), PP2CM (**C**), P-BCKD (**D**), BDK (**E**), LAT1 (**F**), SNAT1 (**G**), and P-S6K1 (**H**) were measured by western blotting in plantaris muscle tissue collected from control (n = 10) and C26 tumor-bearing (n = 10) mice. Protein levels were normalized to Ponceau staining. Data are presented as mean ± SEM. Statistical significance was determined using unpaired t-tests. **p < 0.01, *p < 0.05.

### Soleus

Following the findings in the red gastrocnemius and plantaris muscle, we then wanted to look at the soleus muscle as the soleus muscle is predominantly composed of Type I (slow-twitch) muscle fibers (41,43). This high proportion of oxidative fibers makes the soleus especially well-suited for sustained, low-intensity contractions, playing a critical role in postural control and endurance-based activities such as walking (43). Owing to its metabolic profile and reliance on oxidative phosphorylation, the soleus is thought to be less susceptible to cancer-induced wasting compared to more glycolytic muscles (46,47). However, its high mitochondrial content and dependence on AA metabolism, including BCAAs (48), make it an important tissue for assessing systemic metabolic disruption.

#### Decreased BCAA concentrations and alterations in AA profiles in soleus muscle of tumor-bearing mice

HPLC analysis revealed significantly reduced levels of valine (p < 0.05), isoleucine (p < 0.01), and total BCAAs (p < 0.05) in the soleus muscle of C26 tumor-bearing mice, while leucine showed a trend toward reduction (p = 0.07) (**Figure 14A**). Glutamate levels trended higher in the C26 group (p = 0.09) (Fig 14**B**). Several other AAs were significantly decreased, including aspartate (p < 0.05), threonine, arginine (p < 0.01), and lysine (p < 0.05), while histidine showed a trend toward reduction (p = 0.06). In contrast, glycine, tyrosine (p < 0.05), and phenylalanine (p < 0.01) were significantly elevated (Fig 14**C**). These findings suggest complex AA remodeling in the soleus during tumor progression, with both depletions and accumulations indicating altered protein turnover and systemic redistribution.

**Figure 14:**
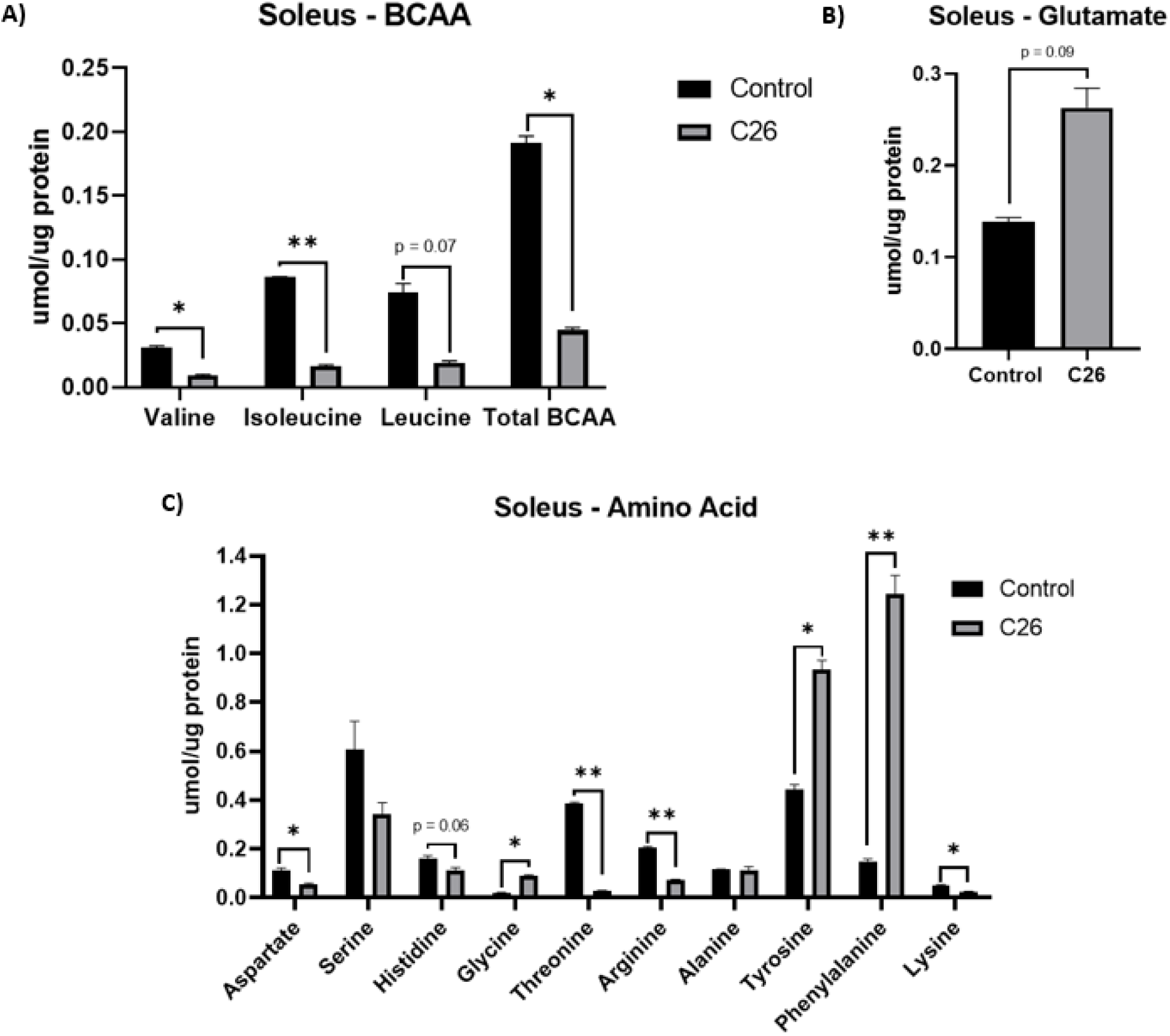
Decreased BCAAs and various alterations in AA concentrations in soleus muscle of 26 tumor-bearing mice. BCAA concentrations (**A**), glutamate (**B**), and other AAs (**C**) were measured in soleus muscle tissue collected from control (n = 5) and C26 tumor-bearing (n = 5) mice. AA concentrations are expressed in µmol per µg of protein. Data are presented as mean ± SEM. Statistical significance was determined using unpaired t-tests. **p < 0.01, *p < 0.05.

#### Reduced LAT1 and P-BCKD expression and elevated P-S6 signaling in soleus muscle of tumor-bearing mice

Due to the small size of the soleus muscle, only a subset of proteins was assessed by western blotting, with limited tissue availability allowing for analysis of P-BCKD, LAT1, and P-S6 (Fig 15**A-D**). Findings revealed significantly decreased P-BCKD expression (p < 0.01) in C26 tumor-bearing mice (Fig 15A-**B**), suggesting increased BCKD enzymatic activity or perhaps alternate post-translation modifications. LAT1 expression was also significantly reduced (p < 0.01) (Fig 15A, **C**), indicating impaired BCAA transport. In contrast, P-S6 was significantly elevated (p < 0.05) in the C26 group (Fig 15A, **D**). These findings suggest that although AA transport may be reduced in the soleus, BCAA catabolism may be upregulated, and anabolic signaling via mTORC1 is elevated, potentially reflecting a compensatory mechanism to mitigate muscle loss or adapt to altered nutrient availability in this highly oxidative muscle.

**Figure 15:**
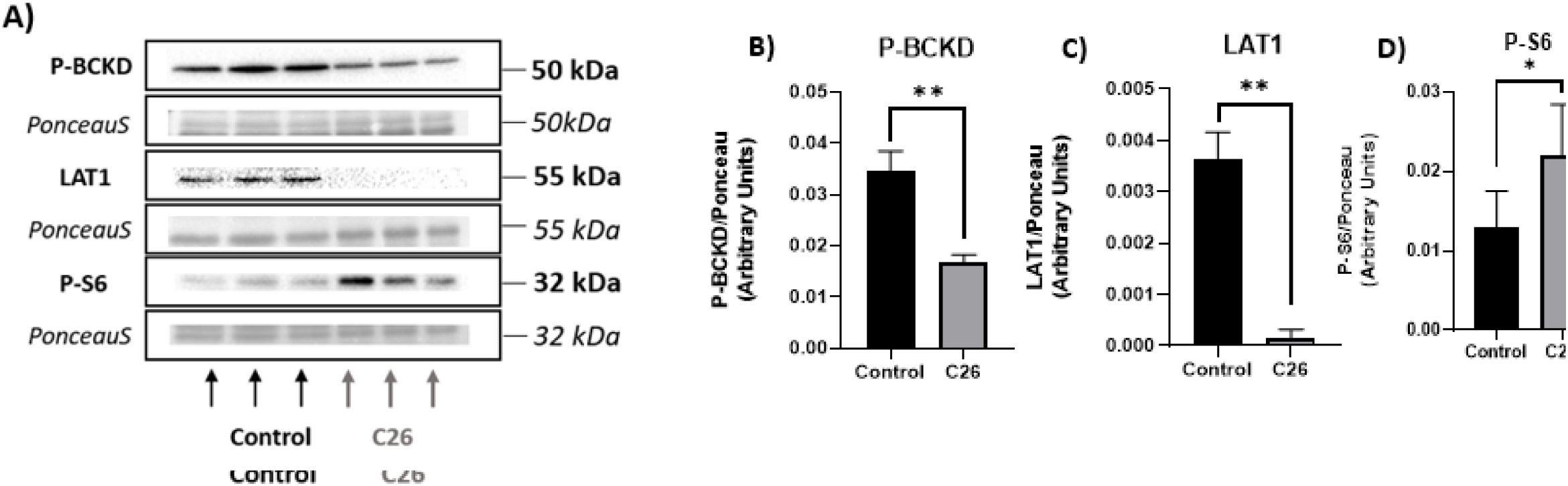
Decreased LAT1, P-BCKD and increased P-S6 expression in soleus muscle of C26 tumor-bearing mice. Representative blots (**A**) and quantified protein expression of P-BCKD (**B**), LAT1 (**C**), and P-S6 (**D**) were measured by western blotting in soleus muscle tissue collected from control (n = 5) and C26 tumor-bearing (n = 5) mice. Protein levels were normalized to Ponceau staining. Data are presented as mean ± SEM. Statistical significance was determined using unpaired t-tests. **p < 0.01, *p < 0.05.

### Overview of Study Findings

*Table 2* provides a comprehensive summary of all major/significant findings across organs and tissues analyzed in this study.

**Table 2:** Comprehensive Summary of Study Findings.

| Tissue | BCAA Levels | BCKA Levels | BCKD Activity | BCKD Regulatory Enzymes | AA Transporters/ mTORC Signaling | Other AA Changes |
| --- | --- | --- | --- | --- | --- | --- |
| Tumor | ↑ | ↔ | ↑ | ↑ PP2CM | ↑ SNAT1, P-S6 | ↔ |
| Liver | ↔ | ↔ | ↔ | ↑ PP2CM, P-BCKD | ↑ P-S6 (p = 0.06)<br>↓ LAT1 | ↑ Lys<br>↓ Asp, Arg, Ala |
| Kidney | ↓ Val | ↓ | ↔ | ↓ PP2CM, P-BCKD | ↑ SNAT1<br>↓ LAT1 | ↑ Arg (p = 0.06)<br>↓ Ala |
| Adipose | ↔ | ↔ | ↔ | ↑ BDK (p = 0.06) | ↓ LAT1, P-S6 | ↑ Ser (p = 0.07), Tyr<br>↓ Glu, Asp, Gly, Thr, Ala, Lys |
| Red Gastroc | ↓ | ↔ | ↓ | ↔ | ↓ LAT1, P-S6 | ↓ Ala, Lys |
| Plantaris | ↓ | N/A | N/A | ↔ | ↓ LAT1 (p = 0.09), ↓ P-S6K1 (p = 0.08) | ↑ Pro |
| Soleus | ↓ | N/A | N/A | ↓ P-BCKD | ↑ P-S6<br>↓ LAT1 | ↑ Glu (p = 0.09), Gly, Tyr, Phe<br>↓ Asp, His (p = 0.06), Thr, Arg, Lys |

↑ indicates increase in C26 group. ↓ indicates decrease in C26 group. ↔ indicates no change between groups. BCAA: Branched-Chain Amino Acids, AA: Amino Acids, BCKA: Branched-Chain Ketoacids, BCKD: Branched-Chain Ketoacid Dehydrogenase, mTORC: Mammalian Target of Rapamycin, Ala: Alanine, Arg: Arginine, Asp: Aspartate, Glu: Glutamate, Gly: Glycine, His: Histidine, Lys: Lysine, Phe: Phenylalanine, Pro: Proline, Ser: Serine, Thr: Threonine, Tyr: Tyrosine

## Discussion

Cancer cachexia is a debilitating condition marked by progressive skeletal muscle wasting, systemic inflammation, and metabolic dysfunction. Central to its pathology is the dysregulation of BCAA metabolism. While the role of BCAA metabolism in tumor cells is well documented (22,23,25), far less attention has been paid to peripheral organs and tissues, such as skeletal muscle, liver, kidney, and adipose tissue. Thus, we investigated the effect of colon cancer (using the C26 tumor model) on BCAA metabolism in the tumor and peripheral sites to examine organ- and tissue-specific metabolic alterations and potential crosstalk between the various tissues in a cancerous environment, a perspective that remains largely undocumented in the field.

We found evidence of coordinated BCAA depletion in peripheral organs and tissues alongside tumor-specific accumulation and upregulation of BCAA transport and catabolism. However, these peripheral sites did not show a uniform compensatory response. While some, like the liver and red gastrocnemius, reduced BCKD activity, possibly as a conservation strategy or due to insufficient BCAA availability to drive BCKD activation, others, such as the kidney and adipose tissue, showed no changes. Transporter expression was also downregulated in several tissues despite low BCAA levels, suggesting impaired uptake or altered regulatory control. Notably, the soleus muscle maintained mTORC1 signaling despite reduced BCAA availability and transporter expression, indicating potential fiber-type-specific adaptations to preserve anabolic signaling.

### Tumor BCAA Accumulation and Systemic Redistribution

One of the main findings of this work was the marked accumulation of BCAAs and the upregulation of AA transporter, SNAT1, and catabolic enzymes in the tumor tissue in week 4 compared to week 2 supporting its role as a metabolic sink, actively drawing in and utilizing circulating AAs to sustain its growth and proliferation [22,23,125]. This aligns with previous studies showing that tumors increase their dependence on EAAs, particularly leucine (49) and glutamine, to support biosynthesis and energy demands (49,50,51). The increased expression of LAT1, a large AA transporter (52,53,54) that is commonly overexpressed in solid tumors (55) including colorectal cancer (56), facilitates this influx of EAAs, stimulating mTORC1 for tumor growth (57), and has been associated with increased tumor aggressiveness and poor prognosis (58,59). Simultaneously, the tumor maintains a highly active BCKD complex, promoting BCAA catabolism to generate substrates for the TCA cycle and lipid synthesis (60). Interestingly, tumor BCKA concentrations remained unchanged, consistent with the idea that elevated BCKD activity efficiently channels available BCAAs into downstream metabolism without substrate accumulation. This strategy is not unique to colon cancer; as similar patterns have been observed in hepatocellular and breast carcinomas, where increased BCKD activity and upregulation of BCAT1 and BCAT2 promote enhanced BCAA catabolism to support tumor growth and proliferation (57,61).

Moreover, these tumor-specific metabolic shifts were paralleled by consistent reductions in BCAA concentrations across peripheral organs and tissues, particularly skeletal muscle (red gastrocnemius, plantaris, soleus) and kidney, suggesting a systemic redistribution of AAs toward the tumor. This pattern aligns with radiotracer studies showing increased leucine uptake in tumors (62,63), reinforcing the concept of nutrient hijacking in cancer cachexia. One possible mechanism contributing to this redistribution is the downregulation of AA transporters, as LAT1 expression was reduced in all peripheral sites examined (liver, kidney, adipose tissue, and skeletal muscle) of tumor-bearing mice. Given LAT1’s role as a bidirectional antiporter facilitating the exchange of large neutral AAs across cell membranes (64,53,54), reduced expression may impair peripheral retention of BCAAs and shift the net flow of these substrates toward circulation, ultimately favoring tumor uptake. This is consistent with prior studies done in our lab, in which chemotherapy-induced cachexia, had also identified LAT1 downregulation as a contributor to reduced intramuscular BCAA levels (20,65), suggesting the idea that transporter regulation may play a central role in cachexia-related AA imbalance, even in the absence of tumor competition (65). Although a lot of literature has shown that LAT1 seems to be upregulated in a vast variety of cancers (including breast, lung and colon among others) (66,67,68,69), the downregulation of LAT1 in peripheral organs and tissues during cachexia appears to be a novel finding and perhaps represents a vulnerability during cachexia progression. In this context, LAT1 downregulation in peripheral sites may not simply be a consequence of tumor burden, but an active contributor to peripheral BCAA depletion and impaired anabolic signaling, one that could present an opportunity for therapeutic intervention.

The loss of peripheral AA levels and their accumulation in the tumor may be further amplified by the tumor-specific upregulation of SNAT1 likely enhancing glutamine uptake to drive LAT1 mediated BCAA import (70,71), supporting continuous nutrient acquisition. This mirrors findings in lung and colorectal cancers, where SNAT1 promotes glutamine metabolism and tumor progression (72). Importantly, glutamine functions not only as a critical exchange partner for LAT1 transport, facilitating the uptake of EAAs like BCAAs, but also as key fuel for nucleotide synthesis, redox balance and the TCA cycle (73). Elevated glutamine levels in week 4 tumors may therefore help sustain mTORC1 signaling and tumor growth (74). This is why glutamine is at the forefront of oncology research and therapeutic interventions have shown promise in targeting tumor progression through glutamine (75,76). Additionally, increased phenylalanine may reflect broader EAA uptake to meet biosynthetic and signaling demands (77), highlighting the tumor’s capacity to reprogram systemic AA metabolism in support of its proliferative needs.

### Liver and Kidney

The next important finding was observed in the organs. Reduction in liver BCKD activity, despite concurrent upregulation of both P-BCKD and PP2CM, suggest a regulatory disconnect that may reflect the liver’s attempt to restrain BCAA catabolism under conflicting metabolic cues. Consistent with this, liver BCKA concentrations remained unchanged, supporting the idea that reduced BCKD activity reflects restrained oxidative flux rather than substrate buildup. This could stem from insufficient substrate availability, as tumor-driven BCAA demand limits systemic supply, or from impaired enzymatic function. In particular, thiamine, a co-factor for the BCKD complex (78), is reduced in gastrointestinal cancers (79,80), and other catabolic conditions such as sepsis and chronic inflammation including diabetes (81), which could potentially reflect reduced BCKD activity in the liver. Alternatively, the BCAA pool may also be preserved in the liver under metabolic stress conditions, such as fasting, trauma, or chronic disease, where skeletal muscle undergoes proteolysis and releases AAs, while the liver conserves key substrates for gluconeogenesis and nitrogen disposal (82). Compared to muscle, which harbors a larger intracellular BCAA reservoir and exhibits greater depletion in this model, the liver may have less capacity but more urgency to retain BCAAs for vital functions consistent with our findings.

Another consideration in interpreting these liver findings is the concept of metabolic zonation within the hepatic lobule. Hepatocytes differ in function depending on their location, with periportal hepatocytes (near the portal triad) exposed to higher oxygen and nutrient levels, and perivenous hepatocytes (surrounding the central vein) operating under relatively lower oxygen and nutrient conditions (83,84). These regions also differ in AA metabolism in which periportal hepatocytes are more active in oxidative pathways such as the urea cycle and gluconeogenesis, whereas perivenous hepatocytes contribute more to glutamine synthesis, glycolysis, and xenobiotic metabolism (84,85). Although BCAA catabolism has not been studied in detail across these zones, it is plausible that BCKD activity and regulation vary between them. This spatial heterogeneity may contribute to variability in hepatic BCKD activity that is not captured by whole-tissue measurements of phosphorylation or protein expression, and could therefore underlie some of the discrepancies observed in our study.

The kidney exhibited more overt signs of AA and BCKA depletion, thus, even as AA concentrations decline, there was unchanged levels of BCKD activity. This pattern suggests reduced systemic availability rather than enhanced renal oxidation. Further, the modest declines in NEAA, such as alanine and arginine, may reflect a selective release of AA that can be endogenously synthesized or are less critical under current metabolic demands. This pattern of selective conservation aligns with findings from fasting, acidosis, and systemic inflammation, where the kidney reduces oxidation of certain AA, particularly BCAAs, while prioritizing glutamine metabolism to sustain ammoniagenesis and acid-base balance (86,87). Further, in models of chronic metabolic acidosis, glutamine uptake by the kidney increases markedly, while leucine and other BCAAs are spared from catabolism (88). This adaptation enhances ammonia production, a key mechanism for neutralizing systemic acidity, and reflects the kidney’s capacity to shift substrate use based on physiological need. Similarly, in the context of cancer-induced stress, the kidney in this study appears to adopt a comparable strategy. Taken together, these findings suggest that the kidney, like the liver, perhaps adopts a restraint based metabolic strategy to preserve core physiological functions in the face of tumor-induced nutrient stress.

Supporting this interpretation, the kidney does not simply filter AAs but actively regulates their reabsorption and metabolism along different nephron segments (89). The proximal tubules are particularly important, as they express neutral AA transporters such as LAT2 (with its heavy chain CD98hc) and SNAT2, which mediate the uptake and exchange of large neutral AAs across the basolateral membrane (90,91). BCAT2 is also expressed in renal tissue and contributes to local BCAA transamination (92). As mentioned earlier, under catabolic conditions these systems adapt by shifting glutamine handling and conserving BCAAs (86,87,88), a process that provides a mechanistic basis for our observation of stable BCKD activity despite overall AA depletion.

### Adipose

Another interesting finding was that although adipose tissue BCAA and BCKA levels remained unchanged, tumor-bearing mice exhibited clear signs of metabolic reprogramming within this tissue, as evidenced by decreases in other AAs, namely glutamate, alanine, aspartate, and glycine, as well as reductions in key metabolic markers LAT1 and P-S6. These changes may contribute to enhanced lipolysis and fat mass depletion, aligning with our data that tumor-bearing mice exhibited significant losses in both body weight and adipose tissue mass (27). Moreover, the reduced availability of glutamate and alanine, which support gluconeogenesis and nitrogen shuttling (93,94), may signal energy stress within adipocytes, activating AMP-activated protein kinase (AMPK), which is a key energy sensor that stimulates lipolysis under nutrient-deprived conditions (35), and promoting lipolytic pathways (95,96). Glutamate also supports mitochondrial respiration and TCA cycle function in adipocytes and its depletion may impair oxidative metabolism, further shifting cells toward mobilizing lipid stores (97). Additionally, reductions in LAT1 and P-S6 may impair protein turnover and mitochondrial integrity (96), accelerating adipose catabolism. Some of these findings are consistent with broader literature describing adipose tissue dysfunction under metabolic stress. For example, alanine export during fasting contributes to hepatic gluconeogenesis (98), indicating a similar catabolic adaptation in cachexia. Additionally, reduced P-S6 signaling observed here mirrors suppressed mTORC1 activity seen in adipose tissue in other catabolic states. For example, in models where S6K1 is genetically ablated or pharmacologically inhibited, adipose tissue exhibits reduced mass and altered lipid handling, closely resembling the cachectic phenotype (99). These findings contrast with obesity, where mTORC1 and P-S6 signaling are elevated to support adipose expansion and lipid synthesis (100), highlighting how nutrient sensing in adipose tissue is tightly linked to systemic energy demands.

Additionally, while reduced caloric intake may contribute to fat loss (101), tumor-derived factors such as zinc-α2-glycoprotein (ZAG) and lipid-mobilizing factor (LMF) directly stimulate lipolysis by activating hormone-sensitive lipase (HSL) and adipose triglyceride lipase (ATGL), independent of food intake (102). Notably, emerging evidence suggests that AA depletion may enhance adipose responsiveness to these signals. For instance, metabolically stressed adipocytes exhibit heightened lipolytic responses especially in tumor microenvironments (103) linking nutrient stress to exaggerated fat mobilization. In this context, AA dysregulation may not simply reflect adipose wasting, but actively potentiate it by impairing mitochondrial metabolism, blunting anabolic signaling, and priming adipose tissue for catabolic activation. Thus, AA metabolism may be intersecting with lipid mobilization as both a target and facilitator of tumor-induced wasting, underscoring its role as a central node in adipose tissue remodeling during cachexia.

### Skeletal Muscle: Fiber-Type Specific Disruptions

In the red gastrocnemius, we observed pronounced reductions in BCAAs, BCKD activity, LAT1 expression, and P-S6 levels, while BCKAs remained unchanged, indicating a strong catabolic shift and impaired AA utilization. This aligns with previous cachexia models showing suppressed leucine signaling even in mitochondria-rich fibers (104,105). The decline in BCKD activity, despite unchanged regulators, may reflect post-translational inhibition, such as impaired cofactor availability (such as thiamine) (106) or reduced substrate flux, mechanisms also implicated in other atrophy models (107). The plantaris showed milder disruption, with partial reductions in BCAAs, LAT1, and P-S6, suggesting intermediate susceptibility.

In contrast, the soleus displayed a distinct and potentially protective profile. Despite reduced BCAAs and LAT1, it maintained muscle mass (27) and exhibited elevated P-S6, consistent with preserved mTORC1 signaling. This aligns with prior studies showing that muscle fiber type strongly influences susceptibility to cachexia, with fast-twitch muscles more prone to atrophy and oxidative, type I fiber-rich muscles more resilient (46,47). In our model, this was reflected by the atrophy of red gastrocnemius and plantaris muscles and preservation of the soleus muscle that we reported (27). Type I fibers are characterized by high mitochondrial density, enhanced oxidative capacity, and elevated expression of PGC-1α, a key regulator of mitochondrial biogenesis and anti-atrophic signaling (108,109). These features enhance energy efficiency and may suppress ROS accumulation and proteolytic signaling (47), while typically showing lower expression of MuRF1 and Atrogin-1 under cachectic conditions (110). The elevated P-S6 in the soleus may therefore reflect a fiber-type–specific adaptive mechanism to preserve anabolic signaling and muscle integrity under metabolic stress. Interestingly, despite its oxidative profile, the red gastrocnemius did not exhibit the same protection, suggesting that oxidative phenotype alone may be insufficient. Other factors, such as fiber subtype composition, local metabolic demands, or muscle-specific function, may also determine susceptibility to tumor-induced atrophy. For instance, tyrosine was significantly increased in the soleus of tumor-bearing mice and this AA, in particular, has been shown to help sustain anabolic signaling by enhancing leucine induced mTORC1 activation (111). This has been demonstrated in vitro and ex vivo, where tyrosine lowered the leucine threshold required for S6K phosphorylation and boosted mTORC1 engagement (112).

Another important finding of this work was the mismatch between BCKD activity and its phosphorylation status across several peripheral sites, suggesting additional regulation beyond the reversible phosphorylation of the E1α subunit by BDK and PP2CM (18,20). Prior studies have shown that BCKD is also subject to allosteric control, whereby its activity can be directly inhibited by reaction products such as NADH and branched-chain acyl-CoAs, even under conditions of low phosphorylation (113). Conversely, BCKAs, particularly KIC, have been identified as potent inhibitors of BDK, thereby reducing phosphorylation pressure and indirectly favoring BCKD activation (114,115). In this context, our measurements revealed tissue-specific patterns of BCKAs: significantly reduced in the kidney, but unchanged in liver, adipose tissue, and red gastrocnemius despite marked BCAA depletion. These differences suggest that BCKAs may contribute differentially to BCKD regulation across tissues. For example, lower renal BCKAs could reduce inhibitory feedback on BDK, reinforcing phosphorylation-mediated restraint of BCKD. In contrast, stable BCKAs in muscle alongside reduced BCKD activity point to other inhibitory mechanisms, such as NADH or acyl-CoA accumulation under cachectic stress, while the preservation of BCKAs in adipose tissue may have counteracted increased BDK expression to help maintain BCKD activity. Taken together, these findings highlight that BCKD function is shaped by the combined influence of phosphorylation status and metabolite availability, as reflected by the tissue-specific BCKA profiles observed in this study. This underscores the importance of considering both regulatory protein expression and allosteric mechanisms when interpreting BCKD activity across peripheral tissues in cancer cachexia

### Limitations

While these findings point to potential direction and future research for intervention, several limitations must be considered when interpreting these results. Firstly, it important to note that while LAT1 and SNAT1 were selected for their established roles in BCAA uptake and mTORC1 signaling, particularly in skeletal muscle and tumor tissue (64,116,117,118), they are not the predominant transporters in all peripheral sites. In the liver, kidney, and adipose tissue, other transporters such as LAT2 (L-type amino acid transporter 2: SLC7A8), SNAT2 (sodium-coupled neutral amino acid transporter 2:SLC38A2), and B AT1 (sodium-dependent neutral amino acid transporter 1:SLC6A19) play more central roles in regulating AA exchange and flux (49,50). Another limitation of this study includes that BCKA concentrations and BCKD enzymatic activity was not measured in the plantaris and soleus muscles due to limited sample availability. As a result, conclusions regarding BCKA levels and BCAA oxidation in these muscles are based solely on transporter and signaling data, which may not fully capture underlying enzymatic flux. This study also exclusively used male mice, limiting the ability to assess sex-based differences in tissue responses to cancer. Since sex hormones are known to influence AA metabolism, muscle wasting, and mTOR signaling (86,119,120), the findings may not be fully generalizable to females. Also, mice were relatively young at the time of sacrifice (10 - 12 weeks old), which may not accurately reflect the typical age-related context of colon cancer in humans. Younger animals are in a more anabolic state (67), potentially influencing how they respond to tumor burden. In addition, the C26 colon cancer model itself presents limitations. Tumors in this model are implanted subcutaneously rather than arising in the colon, which bypasses the intestinal microenvironment and nutrient absorption context characteristic of human colorectal cancer. The model is also non-metastatic (121), whereas colon cancer in patients is frequently metastatic, particularly to the liver (122), and therefore may not fully capture the systemic metabolic disturbances associated with advanced disease. Finally, the C26 model induces highly aggressive cachexia, with rapid muscle and adipose loss (123) that may not accurately reflect the slower and more variable progression observed in patients.

### Reframing Cancer Cachexia

Taken together, the findings from this study point toward a coordinated, tissue-specific disruption in AA metabolism driven by the metabolic demands of the tumor. The tumor acted as a metabolically dominant tissue, showcasing increased transporter expression, elevated BCKD activity, and heightened mTORC1 signaling. Reflecting an increased capacity to import, oxidize, and utilize BCAAs and other AAs for growth and proliferation. Meanwhile, nearly all peripheral organs and tissues, including skeletal muscle, kidney, liver, and adipose, showed consistent trends of transporter downregulation, reduced AA retention, and suppressed mTORC1 signaling. These findings extend prior cachexia research by showing, for the first time to our knowledge, that BCKA concentrations are differentially regulated across peripheral tissues, and that LAT1 expression is downregulated in multiple non-tumor tissues despite systemic BCAA depletion. In some cases, discrepancies between enzyme activity, regulatory protein expression, and BCKA levels further suggest additional layers of control, including allosteric regulation and regional heterogeneity within organs such as the liver and kidney. This pattern suggests that the tumor may be outcompeting host tissues for available nutrients, contributing to systemic metabolic stress, tissue wasting, and disrupted energy balance.

This imbalance helps explain why BCAA-targeted interventions have produced limited results in cachexia treatment (15,124). While leucine and other BCAAs are essential for activating mTORC1 and maintaining muscle protein synthesis, reduced transporter expression and mTORC1 responsiveness in peripheral tissues may prevent these tissues from utilizing supplemented BCAAs effectively. At the same time, the tumor, through increased AA transporters and mTORC1 activity, may be better equipped to take up and benefit from these nutrients, raising concerns that systemic supplementation may end up supporting tumor metabolism more than aiding recovery. Conversely, BCAA restriction, though it may limit tumor fuel, could further starve already compromised tissues. Altogether, these findings emphasize that cachexia is a whole-body response to tumor-driven nutrient high jacking, highlighting the need for targeted, tissue-specific strategies rather than single-agent nutritional interventions, and reinforcing the importance of thinking beyond the tumor itself and toward supporting the host.

## Data Availability

Data will be made available upon reasonable request.

## Acknowledgements

Thank you to the Muscle Health Research Centre at York University for providing access to the HPLC system and to the Paluzzi laboratory for the use of their imaging equipment.

## Author Contributions

The original animal study and tissue collection were conducted by LJD. MZ analyzed the samples and drafted the manuscript. LJD, CGRP and OAJA reviewed data and edited the manuscript. All authors reviewed and approved the final version of the manuscript.

